# Genetic regulation of lncRNA expression in the whole human brain

**DOI:** 10.1101/2024.05.20.594968

**Authors:** Yijie He, Yaqin Tang, Pengcheng Tan, Dongyu Huang, Yongheng Wang, Tong Wen, Lin Huang, Jia Wang, Lizhen Shao, Qinyu Cai, Zhimou Li, Yueyang Wang, Taihang Liu, Zhijie Han

## Abstract

Long non-coding RNAs (lncRNAs) play a key role in the human brain, and genetic variants regulate their expression. Herein, the expression quantitative trait loci (eQTL) of lncRNAs encompassing ten brain regions from 134 individuals was analyzed, and novel variants influencing lncRNA expression (eSNPs) and the respective affected lncRNAs (elncRNAs) were identified. The eSNPs are proximate to their corresponding elncRNAs, enriched in the non-coding genome, and have a high minor allele frequency. The elncRNAs exhibit a high-level and complex pattern of expression. The genetic regulation is more tissue-specific for lncRNAs than for protein-coding genes, with notable differences between cerebrum and cerebellum. However, it shows relatively similar patterns across the cortex regions. Furthermore, we observed a significant enrichment of eSNPs among variants associated with neurological disorders, especially insomnia, and identified insomnia-related lncRNAs involved in immune response functions. Moreover, the present study offers an improved tool for lncRNA quantification, a novel approach for lncRNA function analysis, and a database of lncRNA expression regulation in the human brain. These findings and resources will advance the research on non-coding gene expression regulation in neuroscience.

## Introduction

Long non-coding RNAs (lncRNAs) constitute a class of non-protein-coding RNAs with a length of more than 200 nucleotides. These lncRNAs are highly represented in the human genome and widely involved in various key cellular and biological processes^1^. Previous studies have shown that the expression of lncRNAs has greater intra-individual heterogeneity and higher tissue specificity than protein-coding genes^2, 3^. Importantly, the lncRNAs also exhibit significantly high transcript abundance in the brain compared to the other human tissues and critically regulate the development of the human brain, neuronal regeneration, maintenance of synaptic plasticity, and pathogenesis of central nervous system (CNS) disorders^4–8^. Strikingly, more than 40% of currently known lncRNAs are specifically expressed in different regions of the human brain, according to statistics from the GENCODE (v39)^9^. LncRNADisease (v2) predicted that approximately 90% of the total lncRNAs are associated with at least one CNS disorder, although less than 2% of these have been rigorously confirmed by molecular biology experiments^10^.

Moreover, millions of common single nucleotide polymorphisms (SNPs) in the human genome are considered as major factors contributing to individual differences in behavioral traits and disease susceptibility. The genome-wide association studies (GWAS) have identified a large number of SNPs significantly associated with brain development and CNS disorders^11^. They exhibit marked tissue specificity^12^ and are mainly located in the non-coding regions of the genome (approximately 93%), especially the lncRNA sequences^13–15^. Importantly, most of these non-coding SNPs play key roles in the regulation of lncRNA expression, which is a prerequisite for acting on proximal protein-coding target genes indirectly, further contributing to the pathogenesis of many CNS disorders^16–18^. For example, The A allele of rs1950834 serves as a protective variant against major depressive disorder (MDD). A key feature of the mechanism is that the protective allele reduces chromatin accessibility by disrupting the binding of transcription factor ETV1, consequently decreasing the expression of lncRNA AL121821.1 in neural progenitor cells^19, 20^. The minor alleles of three GWAS-identified risk variants (rs7704909, rs12518194, and rs4307059 localized in the gene-poor 5p14.1 region) are significantly associated with autism spectrum disorder (ASD), synergistically promoting the expression of lncRNA MSNP1AS. This lncRNA is highly enriched in the postmortem cerebral cortex of ASD patients, inhibiting the production of moesin and resulting in the dysregulation of neuronal architecture^21^.

Based on the above findings, we can infer that the regulation of lncRNA expression by genetic variation in the brain is crucial for the vital neuronal functions and the pathogenesis of CNS disorders. The expression quantitative trait loci (eQTL) analysis has been used extensively as a robust multi-omics integration tool for systematic analysis of the genome-wide correlations between altered gene expression and genetic variations in a specific tissue. Previous studies have applied eQTL analysis to investigate the regulation of gene expression in the human brain. However, the studies mainly focused on the protein-coding genes or involved only the single brain tissues. For example, xQTL Serve^22^, BrainCloud^23^, and Myers *et al.*^24^ explored the effect of genetic variants on the expression of protein-coding genes in the prefrontal cortex. Although the samples used in SNCID^25^, BRAINEAC^26^, GTEx^27^, and MetaBrain^28^ include multiple brain tissues, the former two are concerned only with the expression and regulation of protein-coding genes, and the latter two lack targeted investigations on the functional characterization of eQTL lncRNAs and their related CNS disorders.

In the present study, we addressed these issues using large-scale multi-omics data from the UK Brain Expression Consortium (UKBEC), which comprises ten brain regions of European-descent individuals without neurological disorders^26^. Firstly, lncRNA identification and quantification were conducted using an improved microarray probe re-annotation method. Then, a rigorous genome-wide eQTL analysis of the quantified lncRNAs combined with the quality-controlled genotype data was conducted. We also characterized and compared the profiles of the regulation of genetic variants on lncRNA expression across brain regions. Finally, a new approach and trained prediction models based on the eQTL results were developed to identify the potential CNS disorder-related lncRNAs and delineate their functions (**Figure 1**). In addition, we constructed user-friendly web servers to store the lncRNA eQTL results, trained prediction models, and analysis tools.

**Figure 1.**
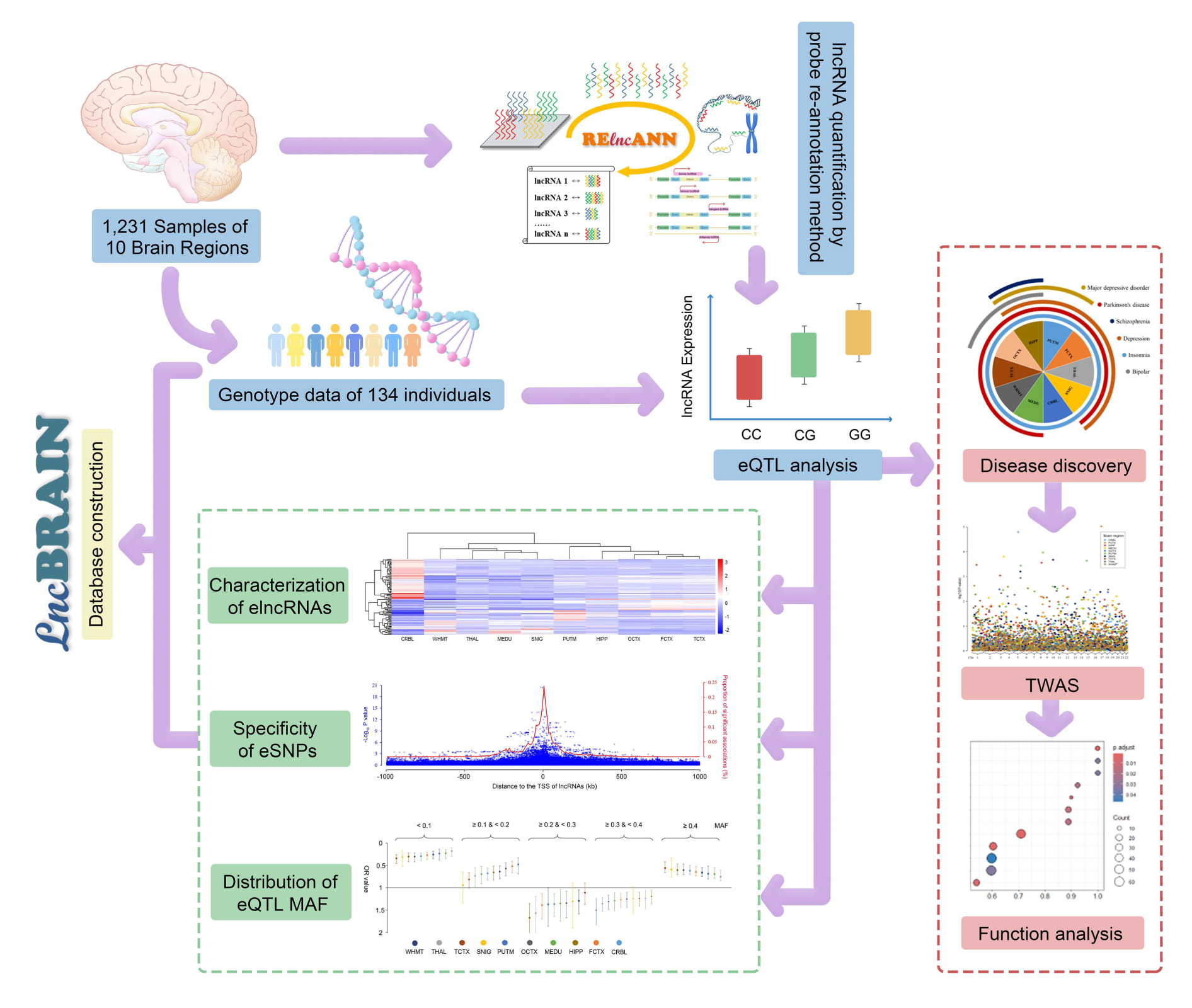
A flowchart of the study to profile the regulation of lncRNA expression in the human brain. The microarray probe re-annotation method was improved to quantify lncRNAs across ten distinct brain regions, utilizing a dataset comprising 1,231 samples sourced from UKBEC. Then, a cis-eQTL analysis of the lncRNAs was performed using the quality-controlled genotype data from the same individuals. Subsequently, the regulation of lncRNA expression across diverse brain regions was comprehensively characterized from multiple perspectives. Finally, a novel approach was designed to identify potential CNS disorder-related lncRNAs and determine their functions, leveraging eQTL results. Moreover, the findings and analytical models of this study were stored in a user-friendly visual database lncBRAIN and the probe re-annotation tool was integrated into a web server RElncANN for future researchers use in neuroscience research.

## Results

### Probe set re-annotation and lncRNA quantification

First, we obtained the 1,231 raw CEL files of Affymetrix Exon 1.0 ST arrays from the Gene Expression Omnibus (GEO) dataset GSE60863. The 1,231 samples from 134 donors of European-descent without neurological disorders involved ten brain regions, i.e., frontal cortex (FCTX), occipital cortex (OCTX), temporal cortex (TCTX), cerebellar cortex (CRBL), hippocampus (HIPP), medulla inferior olivary nucleus (MEDU), substantia nigra (SNIG), putamen at anterior commissure (PUTM), thalamus at lateral geniculate nucleus (THAL), and intralobular white matter (WHMT). The statistics of the samples are summarized in **Methods** and **Supplementary Table S1**.

Then, the pre-labeled exogenous spike-in probes were used to produce quality control metrics of the array using the Expression Console software (v 1.4.1). The signal intensity of the quality control probes shows the expected rank order for all 1,231 samples, i.e., BioB < BioC < BioD < Cre (**Supplementary Figure S1**). Next, we identified and quantified 11,587 lncRNAs (approximately 74.2% of all lncRNAs according to the annotation of Ensembl hg38 release 82) from the1,231 exon arrays using an improved probe set re-annotation strategy (see **Methods** for details). These lncRNAs consisted of 6,222 lincRNAs (53.70%), 4,141 antisense lncRNAs (35.74%), 693 sense intronic lncRNAs (5.89%), 376 processed lncRNAs (3.25%), 154 sense overlapping lncRNA (1.33%), and a macro lncRNA (5.89%) (**Supplementary Table S2**). The proportions of the six identified lncRNA subtypes by re-annotation were consistent with their proportion in the total lncRNAs (50.95%, 36.64%, 6.03%, 5.12%, 1.25%, and 0.01%, respectively) (**Supplementary Figure S2 a** and **b**). The robust multichip average (RMA) algorithm revealed that the overall expression level of these identified lncRNAs was similar across the 10 brain regions (**Supplementary Figure S3**). Finally, we used two metrics, lncRNA recognition rate and average probe coverage, to evaluate the quality of the improved probe set re-annotation approach. The probe coverage of this study was approximately 16.33 probes per lncRNA. Compared to previous re-annotation methods, the lncRNA recognition rate and average probe coverage are about 10.28% and 13.23^29^, 64.37% and 19.83^30^, and 86.94% and 3.35^31^, respectively. According to the results, our method outperformed most of the existing methods in each aspect. Additionally, Zheng *et al.*’s method allows for one-base mismatch during the sequence alignment process^31^. Therefore, our improved probe set re-annotation approach is advantageous in lncRNA discovery and quantification (**Supplementary Figure S2 c**).

### Cis-eQTL analysis of lncRNAs in the ten brain regions

First, the genotype data of the 134 individuals were subject to quality control and their genomic coordinates (hg19) before converting into the GRCh38 human genome (hg38) according to the single nucleotide polymorphism database (dbSNP) annotation (see **Methods** for details); a total of 6,446,207 genotyped variants were obtained for subsequent analyses. Then, the expression data of the same individuals were pooled for a cis-eQTL analysis to assess the genome-wide cis-acting influence of variants on the expression profiles of lncRNAs in the ten brain regions using the R package Matrix eQTL with parameters age, gender, first three genotyping principal components (PCs), and first 15 probabilistic estimation of expression residuals (PEERs) serving as covariates^32^. The cis-window was established within a range of ± 1 Mb around the transcriptional start site (TSS) of lncRNAs. Next, we conducted a permutation procedure to correct for multiple variants in linkage disequilibrium (LD) testing per lncRNA expression and a multiple testing correction to determine the false discovery rates (FDRs) based on Storey approach (the threshold of FDR *q* < 0.05)^33^. Consequently, 440 lncRNAs were identified; their expressions were significantly affected by cis-eQTLs in at least one brain region (elncRNAs). The number of elncRNAs was from a minimum of 70 (SNIG) to a maximum of 172 (CRBL) in the ten brain regions (**Supplementary Table S3**). Finally, we used the first 5% of the permuted minimum *P* values as a threshold to discover the variants with significant regulatory effects (eSNPs) in each lncRNA, as described previously^27, 34^. A total of 25,645 eSNPs were discovered in at least one brain region (from a minimum of 2,367 in SNIG to a maximum of 9,264 in CRBL) (**Supplementary Table S4**). Moreover, a user-friendly lncBRAIN database was constructed to visualize the lncRNA expression, eQTL results, and LD of the corresponding variants in the ten brain regions (http://lncbrain.org.cn/).

### Comparison with the cis-eQTL signals of GTEx

In order to confirm the repeatability of our findings, we evaluated the consistency of the identified elncRNAs and corresponding eSNPs with GTEx datasets (v8) among the five shared brain regions, i.e., CRBL, FCTX, HIPP, SNIG, and PUTM. The overall elncRNAs and eSNPs were identified by pooling the results of the five brain regions. For elncRNAs, we selected 11,320 lncRNAs common to this study and GTEx as the background. **Figure 2a** shows a significant overlap between the elncRNAs identified this study and GTEx based on the hypergeometric distribution test (*P =* 1.38×10^-59^); about a third of the elncRNAs are newly discovered. We also observed similar results in the five brain regions, respectively (**Supplementary Figure S4**). For eSNPs, the SNPs located in cis regions of the overlapped elncRNAs were selected as background. A significant overlap of eSNPs was observed between the two datasets (hypergeometric distribution test *P =* 0) and nearly 40% of our identified eSNPs were novel (**Figure 2a**), which is also consistent with the results of the five brain regions (**Supplementary Figure S5**). In addition, we found that both the recognition rates of elncRNAs and the corresponding eSNPs in different brain regions showed the same trend in our study and that of GTEx (cor = 0.84 and 0.83, *P* = 3.64×10^-2^ and 4.16×10^-2^, respectively) (**Figure 2b**). Next, we compared the number and distribution of the cis-SNPs used for eQTL analysis in this study and GTEx and found that the number of cis-SNPs of the 11,320 common lncRNAs in the two studies have a strong linear correlation (cor = 0.79, *P* = 0) (**Figure 2c**). Nonetheless, the total number of cis-SNPs used in this study is significantly less than that of GTEx (two-tailed Wilcoxon test *P* = 0) (**Figure 2d**), which might be the primary cause of fewer elncRNAs and eSNPs identified in this study compared with GTEx. In summary, the results above reveal a significant replication and general reliability of our findings.

**Figure 2.**
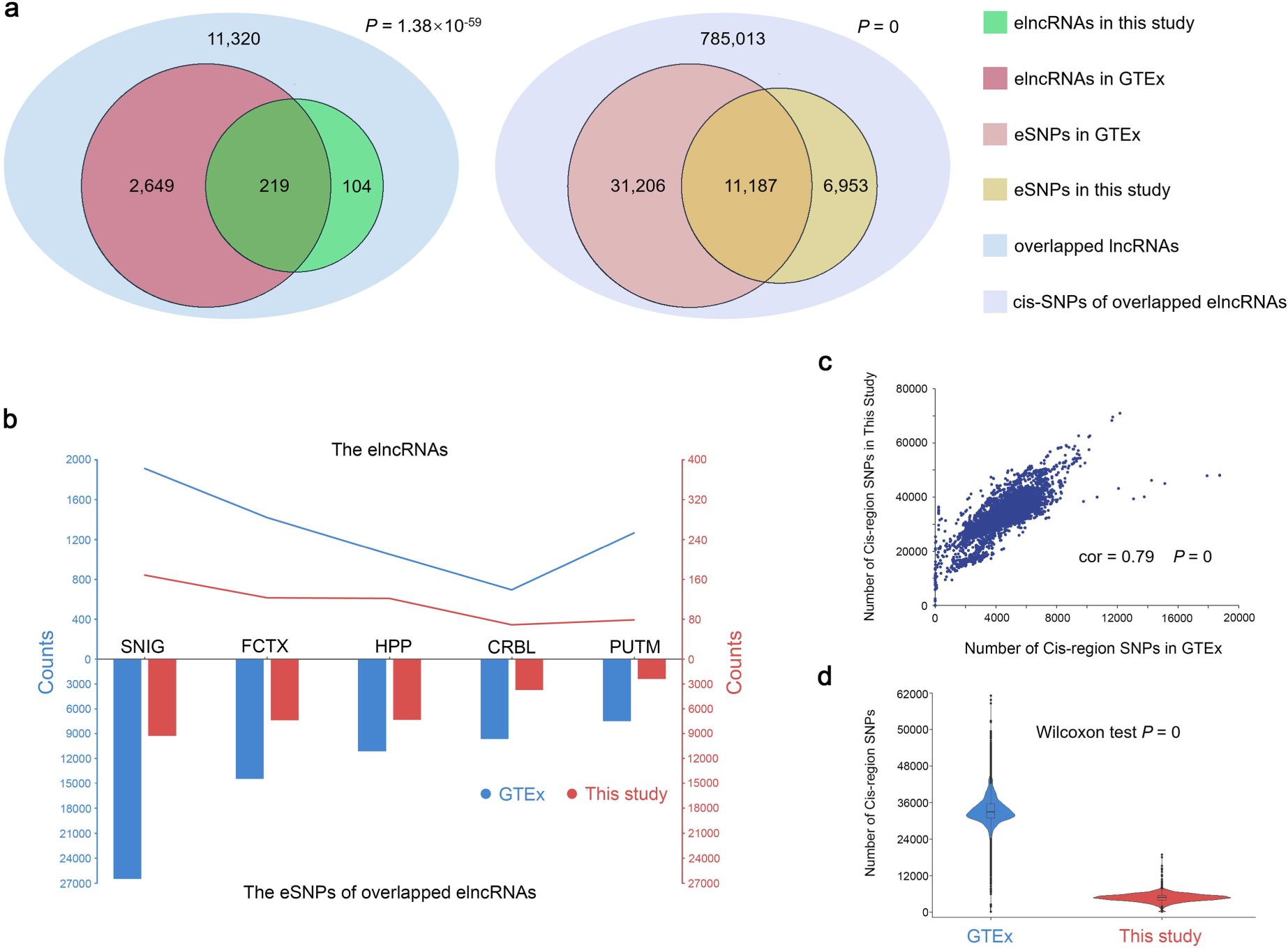
Comparison of the cis-eQTL signals between GTEx and this study. (**a**) The replication of the total elncRNAs (left) and eSNPs of overlapped elncRNAs (right) in the five brain regions (i.e., CRBL, FCTX, HIPP, SNIG, and PUTM) identified by this study in that of GTEx (v8), respectively. The statistical significance was determined by a hypergeometric distribution test with the 11,320 common lncRNAs and the 785,013 cis-SNPs of overlapped elncRNAs as the background, respectively. (**b**) The number of elncRNAs and eSNPs in overlapped elncRNAs in each brain region identified by this study and GTEx, respectively. (**c**) The quantitative correlation of the cis-SNPs between this study and GTEx. Each dot represents the number of SNPs in the cis region of an lncRNA. (**d**) The violin plots show the quantitative distribution of cis-SNPs per lncRNA in the two studies.

### Characterization of the elncRNAs in different brain regions

In order to characterize the elncRNA, we analyzed their expression profiles and features in different brain regions. First, a differential expression analysis was conducted between each brain region and the remaining regions using the R package limma^35^. The significance threshold was set at fold-change (FC) > 1.5 and FDR *q* < 0.05. The two-tailed Fisher’s exact test revealed that the proportion of differentially expressed lncRNAs in the 440 elncRNAs was significantly higher than that in the total 11,587 lncRNAs quantified (odds ratio (OR) = 3.36, *P* = 2.20 ×10^-16^); the results were consistent across the ten brain regions (**Supplementary Table S5**). In addition, elncRNAs showed a marked difference in expression between CRBL and the remaining brain regions, while it was mostly similar among the three cerebral cortex regions (i.e., OCTX, FCTX, and TCTX) (**Figure 3a**). Then, we compared the distributions of the 440 elncRNAs and assessed their similarity among the ten brain regions using the Jaccard distance index (intersection of elncRNAs of two brain regions divided by their concatenation). **Figure 2b** shows a small difference in elncRNA counts among the ten brain regions, although they are most abundant in CRBL. However, a substantially higher number of elncRNAs was detected in CRBL than in the other brain regions. The Jaccard distance index indicated that the similarity is the lowest between CRBL and the other nine brain regions in terms of elncRNAs. It also reveals the highest overall similarity among the three cerebral cortex regions, which is consistent with the findings of the differential expression analysis. Specifically, 119, 125, and 140 elncRNAs were identified in OCTX, FCTX, and TCTX, respectively. About 31.46% elncRNAs were common in the three brain regions, and 48.83% are present in at least two of these regions (**Figure 3c**). Next, we employed a two-tailed Wilcoxon test to assess the differences in characteristic features between the elncRNAs from distinct brain regions; 76 lncRNA features were obtained from the LnCompare database^36^. The results indicated persistent differences in the features, mainly between the CRBL and other brain regions. A total of 39 lncRNA features differed significantly between at least one pair of brain regions (*P* < 0.05), and about 90% of these were involved in CRBL. These differences of lncRNA features were detected with respect to gene/exon length, cell regulation, number of potential risk variants, and distribution in various tissues and cell lines (**Figure 3d** and **Supplementary Table S8**). Finally, we compared the average expression and fluctuations in expression between elncRNAs and non-elncRNAs (empirical *P* > 0.05) in each brain region using the two-tailed Wilcoxon test, respectively. The elncRNAs consistently showed a significantly higher average expression level and variance across individuals than the non-elncRNAs in all ten brain regions (*P* < 0.05) (**Figure 3e** and **Supplementary Figure S6**). This finding was further substantiated by calculating the expression standardized residuals (Z-scores) of the lncRNAs used for eQTL analysis to evaluate their expression levels and fluctuations comprehensively. The Z-scores of each lncRNA in each sample were calculated by regressing the gender, first three PCs, and first 15 PEERs from lncRNA expression levels. The lncRNAs with absolute median Z-score across brain regions greater than two in at least one individual were defined as the outlier (see **Methods** for details)^37^; 322 outliers were detected (**Supplementary Table S9**). The results of the hypergeometric distribution test showed significant overlap (*P* < 0.05) between the elncRNAs and the outliers in each brain region, respectively (**Supplementary Figure S7**). Taken together, the effect of genetic variation on lncRNA expression is specific in the cerebellum and similar between different regions of the cerebral cortex. Interestingly, the lncRNAs with high expression and tissue specificity in brain are likely elncRNAs.

**Figure 3.**
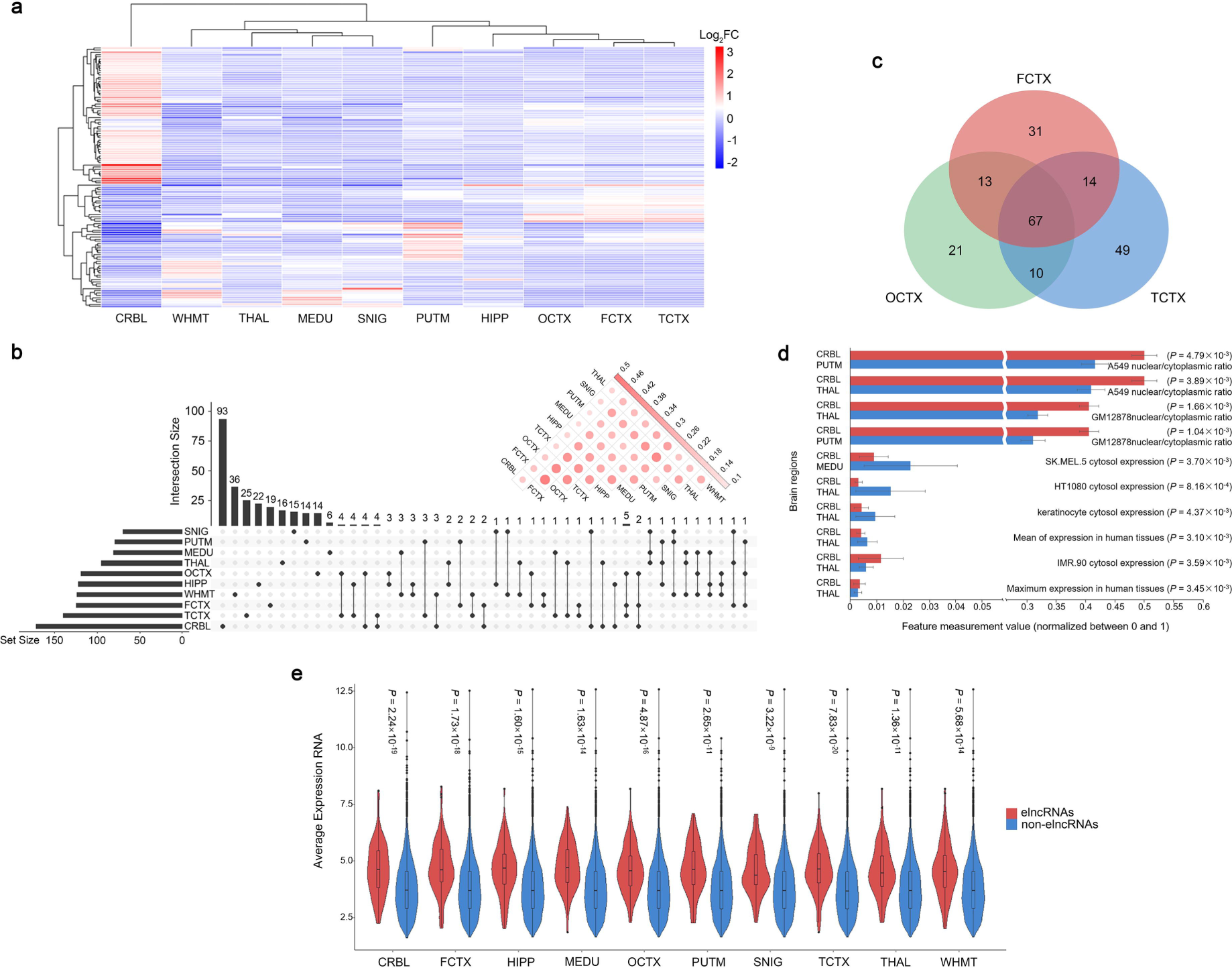
Characteristics of elncRNAs in the human brain. (**a**) Heatmap shows the differential expression pattern of elncRNAs among the ten brain regions. (**b**) UpSet plot shows the number of elncRNAs obtained from different brain regions and their tissue specificity and intersection. Corrplot in the upper right corner reveals the Jaccard distance of elncRNAs among the ten brain regions. The large circles with dark shades indicate high similarity. (**c**) Venn diagram shows the overlap of elncRNAs identified in the cerebral cortex regions, i.e., OCTX, FCTX, and TCTX, respectively. (**d**) The top ten most significant differences in characteristic features between the elncRNAs from distinct brain regions. Each bar represents the average measurement of a feature in a specific brain region. The overall results are summarized in **Supplementary Table S8**. (**e**) Violin plots show the average expression of each elncRNA and non-elncRNA across individuals in the ten brain regions.

### Specificity of genetic regulation in different brain regions

The specificity of genetic regulation on lncRNA expression in different brain regions was assessed by eSNP profiling. Firstly, consistent with the tissue specificity of the elncRNAs, the eSNPs unique to CRBL significantly exceeded those in other brain regions (a similar number of eSNPs in the ten brain regions) (**Supplementary Figure S8**). Then, we explored the association between the effect of variant on lncRNA expression and its distance from the TSS of the corresponding lncRNA. The results showed that the eSNPs tend to be clustered around the TSS of lncRNA, especially for the variants with lower P-values in the cis regions (**Figure 4a**). These findings are consistent across the ten brain regions (**Supplementary Figure S9**). Next, we explored the functional characterization of the eSNPs in ten brain regions. For this reason, the genotyped variants were annotated by ANNOVAR software against the refGene file (hg38)^38^, and the LD-based filtering was carried out using PLINK with the 1000 Genomes Project phase 3 European data (the threshold of r^2^ > 0.8)^39, 40^. The independent variants in cis regions with the eQTL nominal *P* > 0.5 were selected and defined as the non-eQTL SNPs (see **Methods** for details). A total of 2,628 eSNPs and 1,927,132 non-eSNPs were uniquely annotated as one of the 11 functional class (including ncRNA intronic, ncRNA exonic, ncRNA splicing, upstream, downstream, 5’-untranslated region (UTR), 3’-UTR, exonic, intronic, splicing, and intergenic regions) and independent of each other in whole brain. The results showed that the percentage of variants located in non-coding genomic regions (i.e., ncRNA intronic, ncRNA exonic, and ncRNA splicing) is significantly higher in the eSNP set (15.37%, 3.42%, and 0.15%, respectively) compared to the non-eSNP set (6.24%, 0.39%, and 0.002%, respectively) (**Figure 4d**). By comparing the proportions of each functional class between eSNPs and non-eSNPs using the two-tailed Fisher’s exact test with a threshold of *P* < 0.05, we found that the eSNPs are significantly and most highly enriched among variants in ncRNA splicing (OR = 84.00, *P* = 2.72 ×10^-7^), ncRNA exonic (OR = 9.03, *P* = 1.48 ×10^-52^), and ncRNA intronic regions (OR = 2.73, *P* = 2.91 ×10^-61^) (**Figure 4b**). Similar results were observed in each of the ten brain regions (**Supplementary Figure S10**). Furthermore, to assess whether the eSNPs of lncRNAs are enriched in the regulatory elements (e.g., promoter and enhancer) mainly located in the non-coding genomic sequences, ChIP-seq peak data for human transcription factor (TF) binding sites in FCTX from ReMAP (dataset “brain-prefrontal-cortex 2022”, containing two TFs: SIN3A and OLIG2^41^) were used for the two-tailed Fisher’s exact test (see **Methods** for details). The results showed a significant positive enrichment of the eSNPs in both TF SIN3A (OR = 6.17, *P* = 1.54 ×10^-3^) and OLIG2 binding site (OR = 3.71, *P* = 1.42 ×10^-5^) compared with the non-eSNPs (**Figure 4c**). Similar to the results of the previous step, the regulation of lncRNAs by genomic variants mainly involves non-coding functional sequences, which is different from that for protein-coding genes^42^. Finally, we investigated the tissue specificity of genetic regulation on lncRNA expression in different brain regions using an unbiased estimation algorithm for the correlation of beta values (𝑟_𝑏_)^43^. The top-associated eSNPs for each of the elncRNAs in the ten brain tissues were selected to compare the similarity of the genetic effects (measured by beta) between the two tissues, respectively, after correcting for the estimated errors of eQTL analysis (see **Methods** for details). Strikingly, a maximal difference was observed in the genetic effects on lncRNA expression between CRBL and the remaining brain regions (median 𝑟_𝑏_ = 0.33), while it was similar among the three cerebral cortex regions (i.e., OCTX vs. FCTX: 𝑟_𝑏_ = 0.61, TCTX vs. FCTX: 𝑟_𝑏_ = 0.60, OCTX vs. FCTX: 𝑟_𝑏_ = 0.59) (**Figure 4e**). The correlation of genetic effects among different brain regions is consistent with the characteristic of elncRNAs. Moreover, the genetic regulation on lncRNA expression is relatively weakly correlated among brain regions compared to coding genes^43^, indicating a more pronounced tissue specificity of the former. In summary, variants in the proximity of TSS and located in the non-coding genomic regions might affect the expression of lncRNAs in the human brain through tissue-specific regulation.

**Figure 4.**
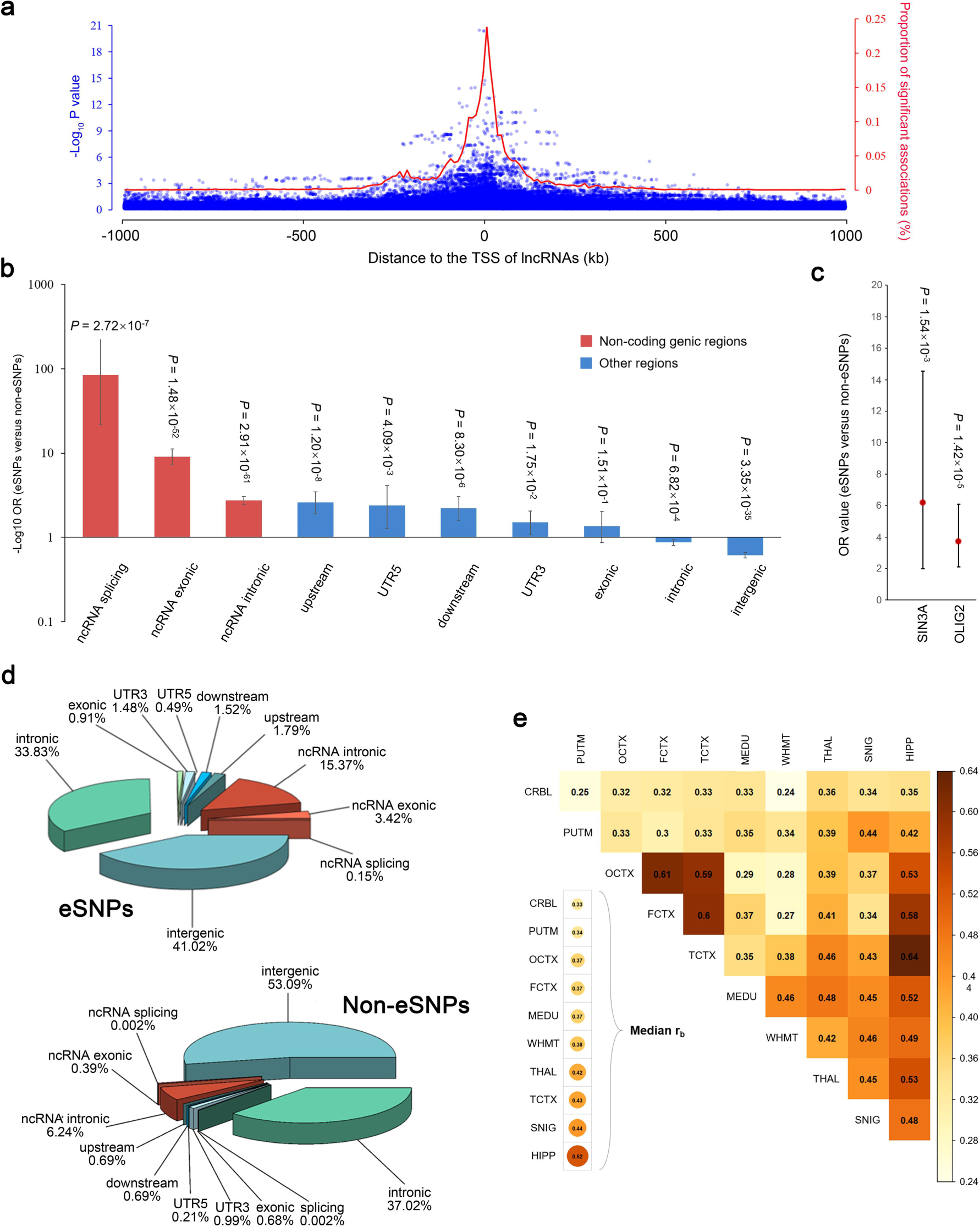
Specificity of genetic regulation on lncRNA expression in the human brain. (**a**) Significance of eQTLs and percentage of eSNPs with respect to the distance from variants to TSS of corresponding lncRNAs in the whole brain. Blue dots represent the variants in cis regions with the negative log-transformed *P* values of eQTL analysis (left y-axis). Red line represents the percentage of eSNPs per 10 kb window (right y-axis). The results of each brain region are displayed in **Supplementary Figure S9**. (**b**) Enrichment analysis of the eSNPs among ten functional categories of variants (i.e., ncRNA splicing, ncRNA intronic, ncRNA exonic, upstream, downstream, 5’-UTR, 3’-UTR, exonic, intronic, and intergenic variants) in the whole brain measured by the two-tailed Fisher’s exact test. The variants in non-coding genomic regions were marked red, while the others were marked blue. The black bars in the histogram indicate 95% CI. The results of each brain region are displayed in **Supplementary Figure S10**. (**c**) Enrichment analysis of the eSNPs among the ChIP-seq peaks of two TFs (i.e., SIN3A and OLIG2) in FCTX by two-tailed Fisher’s exact test similar to subfigure b. (**d**) Pie charts indicate the percentage of each functional type of variants in the datasets of eSNPs and non-eSNPs, respectively. The variants in non-coding genomic regions are shown in warm colors, while the others are shown in cool colors. (**e**) Estimated correlation of genetic regulation on lncRNA expression between two of the ten brain regions. Heatmap of the cells shows the correlation of beta values (𝑟_𝑏_). The median of the 𝑟_𝑏_ between each brain region and the remaining nine areas is displayed in the bottom left corner.

### Correlation of variant frequency and lncRNA cis-eQTL

Herein, we compared the distribution of the LD-pruned eSNPs and non-eSNPs (as defined before) in different minor allele frequency (MAF) bins to elucidate the correlation between the frequency of variants and the magnitude of their effect on lncRNA expression in the human brain. First, we stratified all cis-SNPs into 5% and 10% MAF bins, and extracted the corresponding eSNPs and non-eSNPs from the ten brain regions, respectively. The MAF of variants was calculated based on their genotype dosage values used in this study, and the results are consistent with those of the 1000 Genomes Project phase 3 European population. MAF < 10% criterion was applied as the first bin because the variants with MAF < 5% were removed in quality control filtering (see **Methods** for details). Then, we calculated the proportion of eSNPs and non-eSNPs among cis-SNPs within each MAF bin, respectively. Subsequently, these proportions were evaluated by regression analysis through R function lm in the specific brain region. For the whole brain, the median of the proportion of eSNPs increases significantly with increasing MAF (cor = 0.95, *P* = 3.12×10^-5^) (**Figure 5a**), whereas a significant negative correlation was established between the median of the proportion of non-eSNPs and their MAF (cor = -0.85, *P* = 2.03×10^-3^) (**Figure 5b**). The consistent results were observed in the ten brain regions (**Supplementary Figure S11** and **S12**). Subsequently, we performed a two-tailed Fisher’s exact test to compare the proportion of the LD-pruned eSNPs and non-eSNPs in each MAF bin (see **Methods** for details). Interestingly, the eSNPs were enriched in variants with higher MAF, while the non-eSNPs are more likely to be enriched in variants with lower MAF in the ten brain regions (**Figure 5c** and **Supplementary Figure S13**). Taken together, the altered lncRNA expression profiles in the human brain could be attributed to the regulatory effects of common variants.

**Figure 5.**
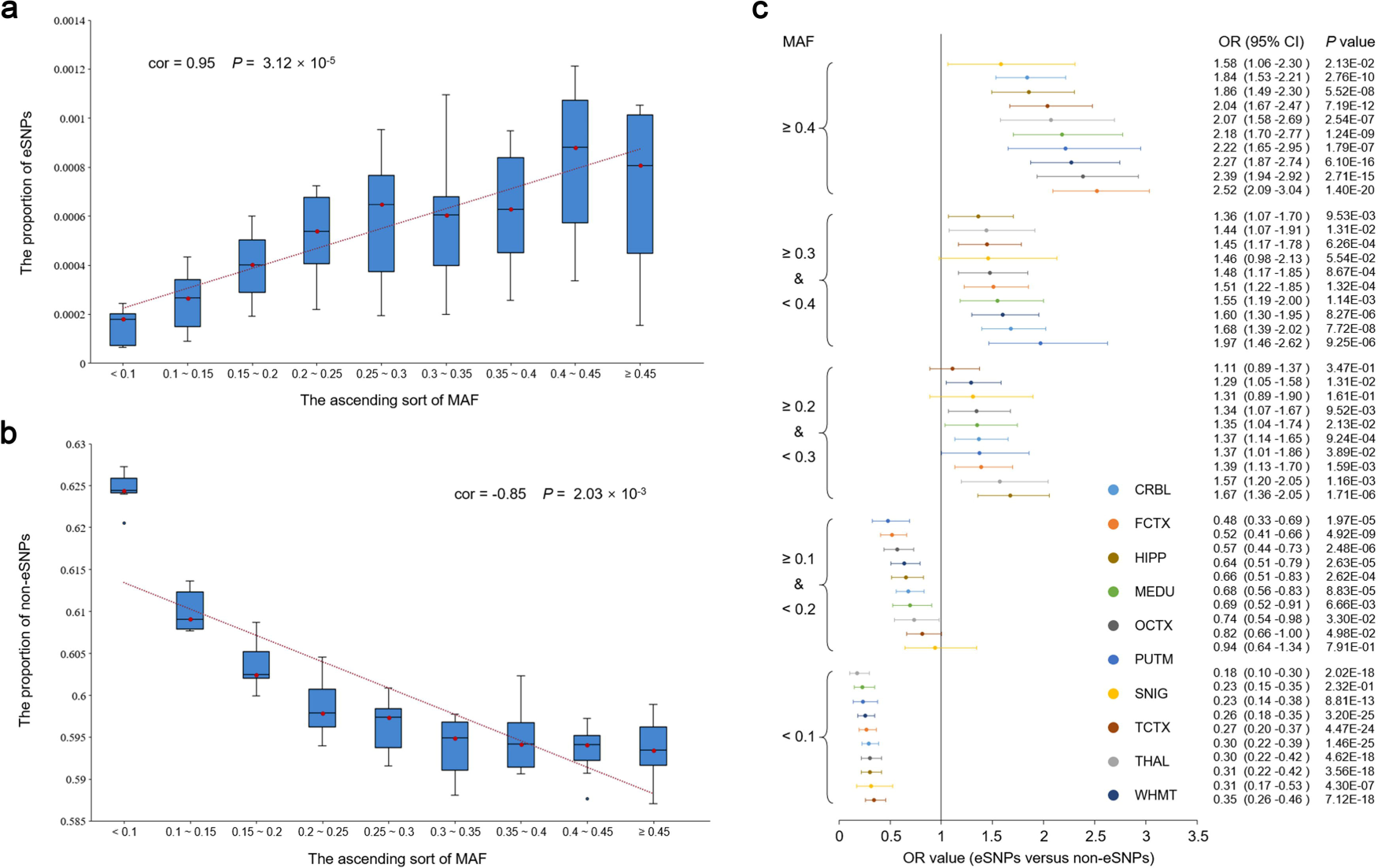
Correlation between variant frequency and their regulatory effect on lncRNA expression in the human brain. (**a**) and (**b**) reveal an increasing and decreasing proportion of eSNPs and non-eSNPs among cis-SNPs with increasing MAF, respectively. The blue box plots show the distribution of these proportions among the ten brain regions in the corresponding MAF bin. The red dotted lines are the linear regression results for the median of the proportions. (**c**) The forest plot shows enrichment results of eSNPs in each different MAF bin by comparing them with the non-eSNPs. The OR and 95% CI were calculated by a two-tailed Fisher’s exact test. Different colors represent different brain regions.

### Disease association of the cis-eQTLs by risk loci enrichment

The basic concept of GWAS is that the common variants tend to contribute to the pathogenesis of common and complex diseases^44^. Given the results of our above analysis and the tissue specificity of eQTL and disease, we speculate that the regulation of lncRNA expression by genetic variability in the brain could be associated with neurological diseases. Thus, we conducted a risk loci enrichment analysis in various human diseases using the association hits (*P* < 1×10^-5^) from GWAS Catalog to assess the potential contribution of the non-coding genetic regulation to disease risks and compare the specificity between neurological and non-neurological diseases^45^. A total of 94 neurological and 417 non-neurological diseases were selected from GWAS Catalog according to the 11th edition of the International Classification of Diseases (ICD-11) (https://icd.who.int/en). Then, the disease-related SNPs were collectively defined as those in LD with the genome-wide significant index SNPs by PLINK based on the 1000 Genomes Project phase 3 European data (the threshold of r^2^ > 0.8)^39, 40^. Each of the 511 diseases was associated with an average of about 1,281 SNPs (**Supplementary Table S10**). Finally, we compared the proportions of the LD-pruned eSNPs and non-eSNPs (as defined before) in each of the disease-related SNPs among the 10 brain regions using the two-tailed Fisher’s exact test (threshold of FDR *q* < 0.05, Benjamini-Hochberg adjustment) (see **Methods** for details). Furthermore, 29 significantly SNP-enriched neurological disease-brain region pairs including six neurological diseases (about 6.4% of all neurological diseases) were identified: insomnia (mean *OR* = 27.73), MDD (mean *OR* = 19.49), Parkinson’s disease (PD) (mean *OR* = 35.07), schizophrenia (mean *OR* = 15.21), depression (mean *OR* = 18.45), and bipolar (mean *OR* = 23.27) (**Figure 6a**). Conversely, only ten such non-neurological disease-brain region pairs (involving about 1.9% non-neurological diseases) showed a close correlation between the lncRNA eSNPs identified in brain tissues and neurological diseases with respect to the diseases (*OR* = 3.47, *P* = 2.82 ×10^-2^) and the disease-brain region pairs (*OR* = 13.23, *P* = 3.15 ×10^-14^) (**Figure 6b**). **Figure 6c** shows that the SNP-enriched neurological diseases encompass more brain regions than the non-neurological diseases. For example, insomnia- and PD-related SNPs were significantly enriched in ten and nine brain regions, respectively, whereas most non-neurological diseases typically involved only one brain region. Moreover, all the six SNP-enriched neurological diseases were identified using HIPP eQTL results, while CRBL was exclusively associated with insomnia. Finally, **Figure 6d** illustrates a higher proportion of significantly SNP-enriched diseases within the category of neurological diseases compared with the non-neurological diseases across most of the ten brain regions, and most significant difference appeared in the HIPP (*P* = 6.87×10^-4^). In summary, these results showed that the influence of genomic variants on lncRNAs might contributes to disease pathogenesis and is mainly associated with neurological diseases; this phenomenon is consistent with the utilization of brain tissue samples in this study.

**Figure 6.**
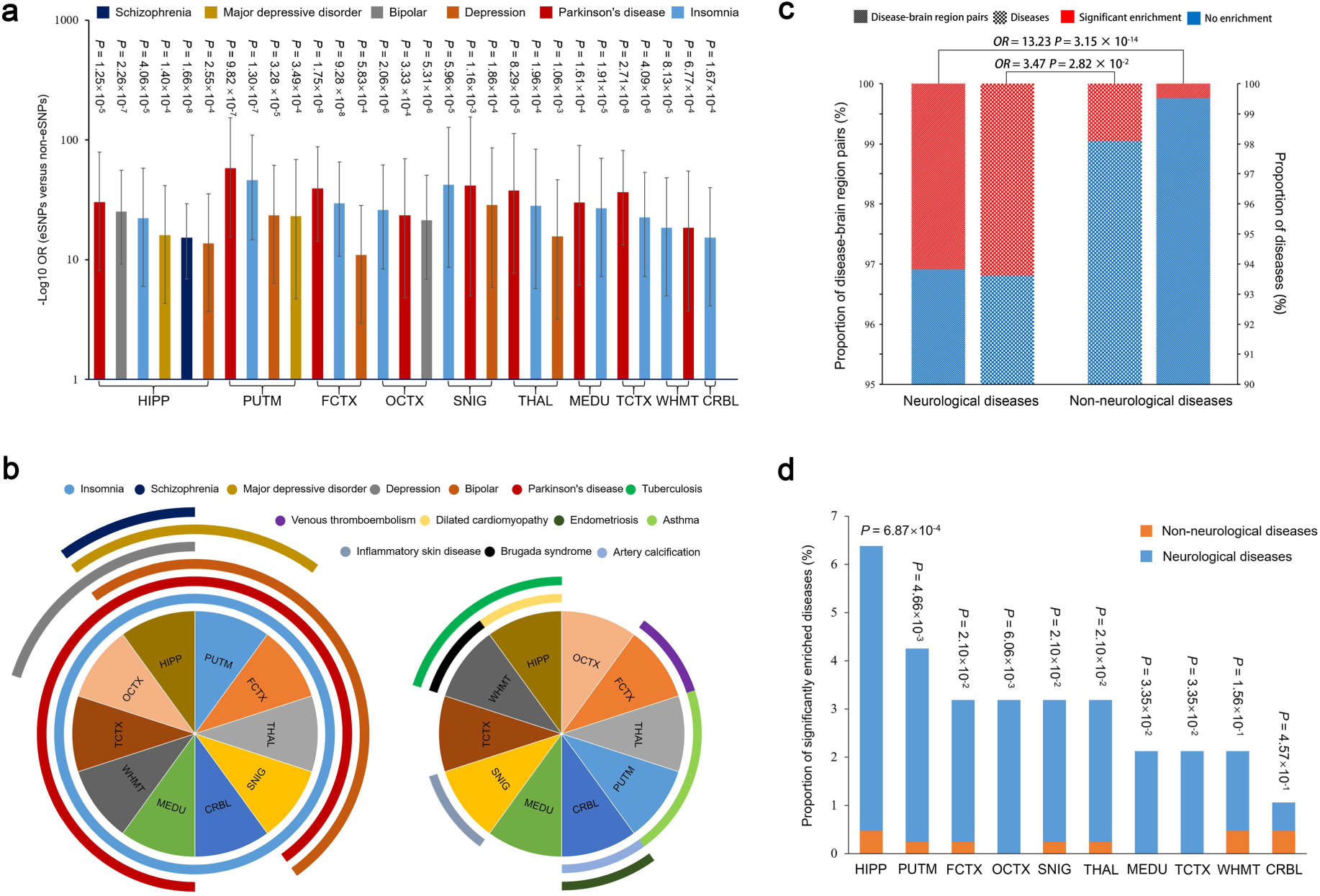
Enrichment of eSNPs among neurological and non-neurological disease-related loci in ten brain regions. (**a**) The 29 significant enrichment results of neurological diseases (i.e., insomnia, MDD, PD, schizophrenia, depression, and bipolar) in the ten brain regions. The two-tailed Fisher’s exact test was employed to explore the enrichment of the eSNPs among 511 diseases (average about 1,281 associated SNPs per disease) compared with non-eSNPs. (**b**) A significantly higher identification rate of SNP-enriched neurological diseases compared with non-neurological diseases. Two-tailed Fisher’s exact test was used to compare the proportion of SNP-enriched neurological and non-neurological diseases, and SNP-enriched neurological disease-brain region pairs and non-neurological disease-brain region pairs in their respective groups. (**c**) The distribution of SNP-enriched neurological and non-neurological diseases in the ten brain regions, respectively. Curve coverage means a significant enrichment of eSNPs among the corresponding disease-related loci in specific brain regions. (**d**) The identification rate of SNP-enriched neurological and non-neurological diseases in the ten brain regions. Two-tailed Fisher’s exact test was used to assess their difference.

### Insomnia-related lncRNAs and their functions

Since only insomnia exhibited significant involvement across all ten brain regions in the risk loci enrichment analysis (**Figure 6c**), we chose insomnia as a representative example to discover the disease-related lncRNAs and investigate their functions based on the eQTL data. Firstly, we utilized eQTL data to train the transcriptome-wide association studies (TWAS) prediction model of lncRNAs using PredictDB, and then combined it with a large-scale GWAS dataset (n = 1,331,010) to identify insomnia-related lncRNAs in the ten brain regions (see **Methods** for details). Subsequently, 169 lncRNAs potentially related to insomnia (*P* < 0.05), including eight with FDR *q* < 0.1 (Benjamini-Hochberg adjustment), were identified (**Figure 7a** and **Supplementary Table S11**). These lncRNAs are implicated in all ten brain regions and do not exhibit any chromosome specificity; however, they are consistent with the elncRNAs in each brain region based on the hypergeometric distribution test (*P* < 0.05) (**Figure 7b**), indicating a critical role of genetic regulation on lncRNA expression in disease mechanisms.

**Figure 7.**
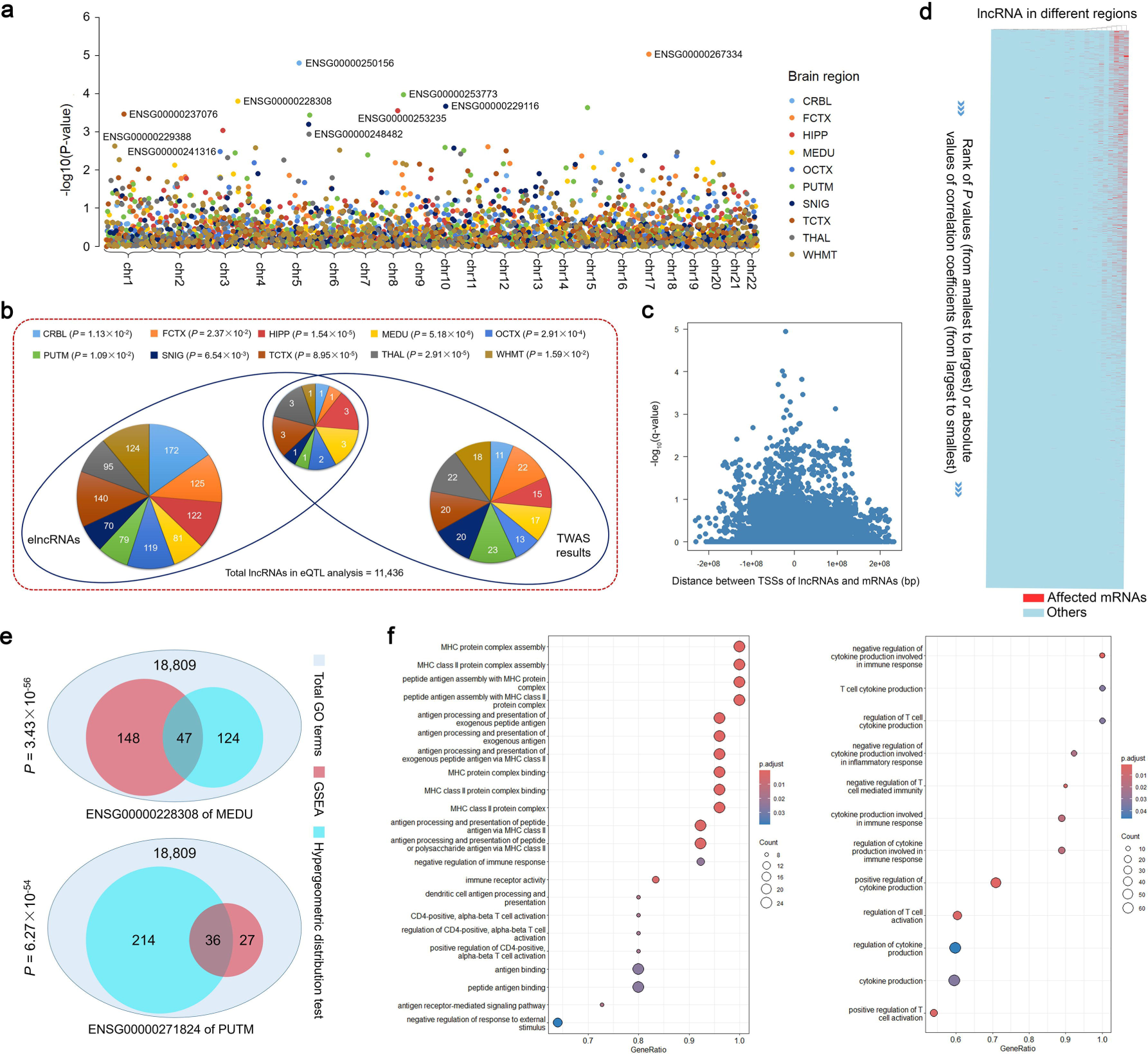
The exploration of insomnia-related lncRNAs and their functions based on eQTL data. (**a**) The Manhattan plot displays the insomnia TWAS outcomes for lncRNAs across ten distinct brain regions. The identified lncRNAs with the smallest *P* value in each brain regions are labeled in the figure. (**b**) The significant consistency between elncRNAs and TWAS results in each brain region was assessed using the hypergeometric distribution test. The distinct brain regions are indicated by different colors. (**c**) The significance of the effect of the lncRNA on mRNA expression is associated with the distance between its TSS when they are located on the same chromosome. (**d**) The heatmap shows the distribution of the mRNAs whose expression is significantly affected by the lncRNAs identified by 2SLS analysis. The x-axis represents the descending order of the absolute values of the correlation coefficients calculated by co-expression analysis between lncRNAs and mRNAs (or ascending order of *P* values). The affected mRNAs (red) are mainly concentrated in high-correlation areas. (**e**) The significant overlap between the lncRNA-related pathways identified by GEAS and hypergeometric algorithm was evaluated using the hypergeometric distribution test. (**f**) The dot plots depict the immune-related pathways (i.e., regulation of cytokine production, T cell-mediated immunity, and antigen processing and presentation) in which two lncRNAs significantly associated with insomnia are implicated.

Since lncRNAs exert their functions by regulating mRNA expression, we designed an approach based on two-stage least squares (2SLS) estimation to discern this causal influence by leveraging eQTL data. The 16,945 protein-coding genes were quantified in the same samples used for lncRNA quantification (see **Methods** for details). In nine brain regions (excluding OCTX), 25 out of 169 lncRNAs, potentially related to insomnia, can significantly influence the expression of at least one mRNA, with a median of about 22 affected mRNAs per lncRNA (the threshold of FDR *q* < 0.05, Benjamini-Hochberg adjustment) (**Supplementary Table S12**). A proximal localization (measured by the distance between their TSSs) of the lncRNA and mRNA on the same chromosome would further enhance the influence effect (**Figure 7c**). Moreover, a co-expression analysis was conducted to calculate the Pearson correlation coefficient of expression between the 25 lncRNAs and 16,945 protein-coding genes pairwise in the corresponding brain regions (**Supplementary Table S13**). Strikingly, the results of 2SLS estimation were consistent with those of co-expression analysis. The majority of the affected mRNAs identified through 2SLS estimation exhibit a pronounced tendency to display stronger co-expression with their corresponding lncRNAs (higher absolute value of Pearson correlation coefficient and smaller *P* values), underscoring the reliability of the 2SLS estimations (**Figure 7d**).

Finally, based on the affected mRNA, we employed two enrichment methods, gene set enrichment analysis (GSEA) and hypergeometric distribution test, to deduce the functions of insomnia-related lncRNAs using the R package clusterProfiler^46^. The Gene Ontology (GO) dataset was used as the reference, and two lncRNAs, ENSG00000228308 (in MEDU) and ENSG00000271824 (in PUTM), were selected according to the following criteria: TWAS *q* < 0.1 and number of affected mRNAs > 20. Interestingly, the enriched terms are mainly involved in immune-related pathways, including regulation of cytokine production, T cell-mediated immunity, and antigen processing and presentation (FDR threshold *q* < 0.05, Benjamini-Hochberg adjustment) (**Figure 7e** and **Supplementary Table S14** to **S15**). Previous studies demonstrated that the human brain contains immune signaling molecules that interact with the neurochemical systems and regulate normal sleep patterns^47^; for example, cytokines have been widely known to play a crucial role in regulating sleep-wake behavior^47, 48^. Several genetic variants associated with sleep duration have been identified within the genes involved in cytokine signaling^49^, and antagonizing cytokines (such as interleukin-1 (IL-1) and tumor necrosis factor (TNF) in human CNS, as well as upd2 in *Drosophila*) can increase the risk of insomnia through endocrine and inflammatory regulation^47, 50^. On the other hand, a negative feedback mechanism was discovered wherein sleep deprivation triggers the efflux of prostaglandin D2 across the blood-brain-barrier, leading to a cytokine storm-like syndrome and severe inflammation in the peripheral immune system^51, 52^. Some studies also noted the interaction between the CD4^+^ T cells and sleep behavior disorders^53, 54^. Therefore, dysregulation of these lncRNAs could be a potential contributing factor to the aforementioned mechanisms of insomnia.

## Discussion

LncRNAs are prevalent in the human genome and intricately involved in diverse cellular and biological processes^1^. Unlike protein-coding genes, lncRNAs exhibit greater intra-individual heterogeneity and tissue specificity^2, 3^. Particularly enriched in the human brain, lncRNAs are key regulators in brain development, neuronal regeneration, synaptic plasticity maintenance, and CNS disorder pathogenesis^4–8^. Many of the variants associated with brain-related traits identified by GWAS are localized in the non-coding genome and regulate lncRNA expression^11–18^. Understanding the genetic regulation of lncRNAs in the brain is crucial for deciphering neuronal functions and CNS disorder mechanisms. Most eQTL analyses focus on protein-coding genes or single brain tissues, lacking in-depth investigations into the functional characterization of eQTL lncRNAs and their contributions to CNS disorders across the brain^22–28^.

In the present study, we integrated eQTL analysis with non-coding gene research to gain an insight into the genome-wide cis-action of variants on lncRNA expression throughout the ten brain regions and highlighted its significance in human CNS disorders. We applied an enhanced probe set re-annotation strategy to quantify 11,587 lncRNAs in 1,231 samples from 134 European descent individuals without neurological disorders and conducted the cis-eQTL analysis in combination with the genotype data of 6,325,551 variants from the same individuals. Comparison with the GTEx dataset validated our findings and identified novel elncRNAs and eSNPs. As shown in the above results, the elncRNAs exhibit higher expression levels and more complex expression patterns than the non-elncRNAs that are unlikely to be regulated by genomic variants across the brain. On the other hand, eSNPs have distinct characteristics compared to the non-eSNPs that do not affect lncRNA expression: (1) shorter distance to the TSS of the corresponding lncRNAs, (2) stronger enrichment among the variants located in non-coding and regulatory element genomic regions, and (3) higher MAF, across the whole brain. Moreover, the genetic regulation of lncRNA expression demonstrated greater tissue specificity compared to that of coding genes. The potency of genetic regulation on expression and the distribution and feature of elncRNAs exhibit differ markedly between the cerebrum and cerebellum while displaying the highest similarity among the cerebral cortex regions. Additionally, we found that genomic variants affecting lncRNA expression in the brain play a critical role in disease pathogenesis, especially neurological disorders. A novel approach based on 2SLS algorithm was developed to explore the biological functions of the predicted insomnia-related lncRNAs. Also, we identified the immune-related pathways that could elucidate aspects of the mechanism underlying insomnia.

In conclusion, this study systematically analyzed the regulation of lncRNA expression based on genomic variants in the whole human brain. It offers valuable insights into the functional characterization of elncRNAs, their corresponding eSNPs, and their significance in human CNS disorders. In this process, we offered the following: (1) an improved probe set re-annotation method, enabling efficient utilization of the extensive microarray data resources already available; (2) a novel approach for the functional explanation of lncRNAs; (3) a database for storage and display of lncRNA expression regulation data in the human brain. These findings and methodologies serve as valuable resources for advancing the research on the regulation of non-coding gene expression in neuroscience.

## Methods

### Genotype and microarray data from ten brain regions of 134 individuals

A total of 1,231 samples (ten brain tissues) of 134 European-descent individuals free of neurological disorders from UKBEC were used in this study^26^. Briefly, the postmortem brains were selected from two sources: the Medical Research Council Sudden Death Brain of Edinburgh and the Sun Health Research Institute Brain Bank of USA. The average age of these individuals at the time of death was about 58 years (ranging from 16 to 102), the mean sample size per brain region was about 123 (ranging from 101 to 131), and the sex ratio (female to male) of these individuals is about 0.35 (**Supplementary Table S1**). Then, for transcriptome profiling, total RNA was extracted from each brain tissue using a single-step RNA isolation method and the miRNeasy 96 kit. After the quality control checks on the Agilent 2100 Bioanalyzer and RNA 6000 Nano Kit, 1,231 qualified samples were processed on 96-well plates at microarray facility and hybridized to Affymetrix Exon 1.0 ST arrays. Subsequently, the arrays were scanned and inspected visually using Affymetrix GeneChip Scanner 3000 7G. The Affymetrix Exon 1.0 ST array CEL files were stored in the GEO database with the accession code GSE60863. Finally, for genotype profiling, genomic DNA was isolated from each brain tissue using DNeasy Blood & Tissue Kit (Qiagen) and subjected to genotyping using Illumina Infinium Omni1-Quad BeadChip and Immunochip. Then, the array scan was carried out on an Illumina iScan system, followed by SNP calling and imputation using Illumina GenomeStudio and MaCH/minimac, respectively. The variants with MAF < 5% and low-quality genotype data were removed. The filtering criteria included the samples with < 95% call rate, suspected non-European ancestry or cryptic sex status and relatedness, variants with false allele or heterozygote, missing genomic location info, genotyping call rate < 95% or Hardy-Weinberg equilibrium (HWE) P value < 0.0001. The more detailed description of sample and upstream analysis is provided in Ramasamy *et al.*^26^. Furthermore, we converted the genomic coordinates of these variants (hg19) into the GRCh38 human genome (hg38) according to the annotation in dbSNP (version 151, human 9606)^55^. Finally, the mismatched variants were removed to obtain a total of 6,446,207 genotyped variants (hg38) were obtained for the following analyses.

### Probe set re-annotation and lncRNA quantification

The previous probe set re-annotation methods were improved to identify and quantify lncRNAs from the Affymetrix Exon 1.0 ST arrays^29–31^. Specifically, the Ensembl gene annotation file with the corresponding reference sequence (Homo sapiens GRCh38/hg38, release 82) was used for the probe set re-annotation and lncRNA identification. It includes 15,618 lncRNAs, 22,017 protein-coding genes, 28,406 pseudogenes and other non-coding genes^56^. After removing the quality control probes, the remaining 5,383,654 probes were mapped to the reference sequence with no mismatch using SeqMap software^57^. Based on the Ensembl sequence annotations, we removed the two types of unsuitable probes: (1) those mapping to the protein-coding genes, pseudogenes, or other non-coding genes; (2) those multi-mapping to more than one gene, whereas those mapping to multiple transcripts belonging to same gene were retained. According to the principle of matching at least four probes per lncRNA, we identified 11,587 lncRNAs and 189,164 corresponding eligible probes. Next, array quality control was performed using the Expression Console software (v 1.4.1) based on quality control probes. The array hybridization and overall signal quality metrics were determined using the pre-labeled exogenous probes with an increasing hybridization concentration, i.e., the BioB, BioC, BioD and Cre signals are arranged in order from lowest to highest (**Supplementary Figure S1**). Finally, based on the lncRNA probe set re-annotation file, the basicRMA function of R package oligo with default parameters was utilized to normalize raw background intensity of array probes via the RMA algorithm and to quantify the expression level of these identified lncRNAs in the 1,231 samples^58^. We also provided an online probe re-annotation tool RElncANN for identifying and quantifying human lncRNAs from the multi-platform microarray data (https://relncann.cqmu.edu.cn/appcel/).

### Genome-wide cis-eQTL analysis of lncRNAs

First, we removed 121 lncRNAs, not annotated with genomic coordinates by Ensembl (GRCh38, release 82), from the 11,587 quantified lncRNAs (**Supplementary Table S2**). Then, the genomic locations of all the 6,325,551 genotyped variants were mapped to a 1 Mb region upstream and downstream of the remaining 11,466 lncRNA TSSs (cis regions) based on the dbSNP annotation (version 151, human 9606)^55^. After removing 30 lncRNAs that did not contain a variant in the cis regions, we selected 6,325,551 variants within 1 Mb around the TSS of 11,436 lncRNAs. Subsequently, a cis-eQTL analysis was conducted in the ten brain regions using the R package Matrix eQTL, respectively, based on the RMA values of the 11,436 lncRNAs in combination with the genotype data of the 6,325,551 variants^32^. A linear regression model was used in the eQTL analysis with parameters age, gender, first three PCs, and first 15 PEERs serving as covariates. Specifically, we used the “transpose trick” in principal component analysis (PCA) to generate the first three PCs of genotype data (high dimensionality). This approach could circumvent a large memory footprint by directly computing the covariance matrix without forming the transpose of the centered data matrix. Factor analysis and Bayesian regression modules were applied to generate the first 15 PEERs from the lncRNA expression data using the R package PEER with parameters age and gender serving as covariates^59^. Subsequently, a permutation procedure was employed to determine the elncRNAs without multiple hypothesis effects of variants in LD, which were used as a basis for the subsequent eSNP discovery, as described previously^27, 34^. Briefly, the minimum nominal *P* value corresponds to the most significant result of cis-eQTL analysis per lncRNA. The permuted *P* values were calculated by randomizing the sample labels of lncRNA expression data and their related covariates (i.e., age, gender, and PEERs), while holding fixed the genotype data and corresponding covariate PCs. The permutations were conducted from a minimum of 1,000 to a maximum of 10,000 and exited when at least 15 minimum permuted *P* values were less than the minimum nominal *P* values. The empirical *P* value of per lncRNA was defined as the ratio of occurrences of extreme events (minimum permuted *P* < minimum nominal *P*) to the total number of permutations (Np). The multiple testing corrections of the empirical *P* value were performed using Storey approach, and the threshold of significant elncRNAs was set at FDR *q* < 0.05. Finally, a permutation threshold of each lncRNA was used to identify the eSNPs. The empirical *P* value of lncRNA whose FDR *q* value was closest to 0.05 (due to the nonlinear transformation of Storey approach) was selected to generate a percentile (*P* × N_p_ of a specific lncRNA), and the permuted *P* value at this percentile (ascending order and ceiling integer) was defined as the permutation threshold for each lncRNA in each brain region. An eSNP was determined if its nominal *P* value was less than the threshold value.

### Differential expression analysis

The differential expression of elncRNAs and total quantified lncRNAs between each brain region and the remaining regions was analyzed, respectively. Firstly, the lmFit function of the R package limma was used to fit a linear model of the expression matrix. Then, the estimated t-statistic and FCs of lncRNAs between each brain region and the remaining regions were calculated by the contrasts.fit function based on the fitted model. Next, the eBayes function was applied to moderate the FCs and to generate the Bayes factors. Finally, the Benjamini-Hochberg adjustment was used for multiple tests based on the ranking of Bayes factors. All functions were executed using the default parameter settings. According to the significance threshold of FC > 1.5 and FDR *q* < 0.05, we identified 116 elncRNAs (**Supplementary Table S6**) and 908 lncRNAs (**Supplementary Table S7**) with differential expression.

### Wilcoxon test

Herein, we used the two-tailed Wilcoxon test to assess the differences in lncRNA expression, characteristic features, and the number of cis-SNPs. Wilcoxon test is a non-parametric statistical test appropriate for the normal or non-normal distribution data. Specifically, a comprehensive feature set of lncRNAs based on the human GENCODE annotation (v24) is cataloged in the LnCompare database, which includes lncRNA length, classification, localization, sequence conservation, nucleotide composition, and transcriptional level in specific cell lines and tissues^36^. Firstly, these features were normalized from 0 to 1 using the R package scales (https://CRAN.R-project.org/package=scales), respectively. Then, the distribution of each normalized feature of elncRNA sets was compared between the pairs of brain regions using the R function wilcox.test with the default parameters. The lncRNAs with empirical *P* > 0.05 were defined as the non-elncRNAs, and the distribution of their average expression levels and variances were compared across individuals per brain region with the elncRNAs by the two-tailed Wilcoxon test, respectively. Finally, we selected the SNPs located in the cis region of the lncRNAs common to this study and GTEx (used for the eQTL analysis) from the two studies, respectively, and compared their number using the two-tailed Wilcoxon test. The significance threshold of the Wilcoxon test was set at *P* < 0.05.

### Fisher’s exact test

The two-tailed Fisher’s exact test was conducted with a 2 × 2 table in this study for the elncRNA characterization analysis (rows of 2 × 2 table: differentially expressed and non-differentially expressed lncRNAs, columns: elncRNAs and total lncRNAs), type enrichment analysis of eSNPs (row: SNPs within and not within a type, columns: eSNPs and non-eSNPs), TF binding site analysis (row: SNPs within and not within regions of TF peaks, columns: eSNPs and non-eSNPs), comparison of variant frequency (row: SNPs within and not within a MAF bin, columns: eSNPs and non-eSNPs), disease association analysis (rows: disease- and non-disease-related SNPs, columns: eSNPs and non-eSNPs), comparison of SNP-enriched diseases (rows: proportion of SNP-enriched neurological and non-neurological diseases, columns: total neurological and non-neurological diseases), and disease-brain region pairs (rows: proportion of SNP-enriched neurological and non-neurological disease-brain region pairs, columns: total neurological and non-neurological disease-brain region pairs), respectively. These analyses were carried out using R function fisher.test.

### Hypergeometric distribution test

The hypergeometric distribution test was applied to demonstrate the overlap between the different sets using the R function phyper(k, m, N-m, n, lower.tail = FALSE). In the overlap of elncRNAs identified by GTEx and this study, N indicates the 11,320 lncRNAs shared by the eQTL analysis of GTEx and this study, m refers to the number of elncRNAs identified by GTEx, n indicates the number of elncRNAs identified by this study, and k refers to the number of elncRNAs shared in GTEx and this study from the corresponding brain regions. In the overlap of eSNPs identified by GTEx and this study, N refers to the number of shared SNPs used for cis-eQTL analysis of the overlapped elncRNAs in GTEx and this study, m indicates the number of the eSNPs identified by GTEx, n stands for the number of eSNPs identified in this study, and k refers to the number of the eSNPs shared in GTEx and this study from the corresponding brain regions. In the overlap of elncRNAs and outliers, N refers to the total of 11,436 lncRNAs used for eQTL analysis and outlier discovery, m indicates the number of elncRNAs, n stands for the number of outliers, and k refers to the number of elncRNAs in outliers from the corresponding brain regions. In the overlap of elncRNAs and TWAS results, N refers to the total 11,436 lncRNAs used for eQTL analysis and TWAS analysis, m indicates the number of elncRNAs, n refers to the number of TWAS findings, and k stands for the number of elncRNAs in TWAS findings from the corresponding brain regions. The significance threshold of the hypergeometric distribution test was set at *P* < 0.05.

### LncRNA expression outlier discovery

The lncRNA expression outliers were discovered using the protocol described previously^37^. Briefly, the standardized expression values of the 11,436 lncRNAs used for eQTL analysis were log2-transformed (i.e., log_2_(RMA+2)) before calculating their mean expression level in each brain region. Then, the mean expression was used as the constant term to regress out the gender, first three PCs, and top 15 PEERs from the corresponding lncRNA expression levels via a linear model using R function lm in each brain region. Furthermore, based on the assumption of normality of the residual vector, we standardized the expression residuals of each lncRNA and yielded their Z-scores in each sample using R function rstandard. Finally, the median of absolute Z-scores for each lncRNA across brain regions for each individual (the individuals with fewer than 5 brain regions were removed) was calculated, and the lncRNAs with absolute median Z-score > 2 at least in one individual were defined as the outlier.

### Generation of independent eSNPs and non-eSNPs as well as disease-related SNPs

We identified both eSNPs and non-eSNPs in the cis regions of lncRNAs. The eSNPs were defined according to their nominal *P* value less than the permutation threshold of the corresponding lncRNAs (as described above). On the contrary, the non-eSNPs were defined based on their nominal *P* > 0.05 (much greater than the permutation threshold) for all lncRNAs, which are unlikely to be associated with their expression in the corresponding brain regions. Then, we performed an LD-based filtering procedure to remove the variants genetically dependent on each other from the eSNPs and non-eSNPs, respectively. Specifically, the *P* value-informed clumping was used to conduct the LD pruning by PLINK based on the 1000 Genomes Project phase 3 European data with the following parameters: window size of 250 kb and r^2^ threshold of 0.8. The number of independent eSNPs and non-eSNPs in the ten brain regions were 647 and 592,436 (CRBL), 548 and 592,683 (FCTX), 471 and 595,568 (HIPP), 345 and 597,597 (MEDU), 461 and 593,728 (OCTX), 245 and 597,813 (PUTM), 167 and 597,099 (SNIG), 544 and 594,139 (TCTX), 295 and 596,337 (THAL), and 550 and 590,628 (WHMT), respectively. Moreover, their numbers in the whole brain were 2629 (concatenation of eSNPs of the ten brain regions) and 1,927,671 (concatenation of non-eSNPs of the ten brain regions after removing eSNPs), respectively.

In order to generate multiple sets of disease-related SNPs, we first obtained the genome-wide significant index SNPs from the GWAS Catalog database according to the threshold of *P* < 1.0×10^-5^. GWAS Catalog provides summary results and comprehensive information about 276,696 index SNPs of 5,481 human diseases and traits^45^. ICD-11 is an international standard classification system of disorders (https://icd.who.int/en) comprising 26 chapters, each corresponding to a specific category of diseases^60^. Chapter 6 (mental behavioural or neurodevelopmental disorders), chapter 7 (sleep-wake disorders), and chapter 8 (diseases of the nervous system) were classified as the neurological diseases, and the remaining were termed non-neurological diseases. Of these, 511 (94 neurological and 417 non-neurological diseases) were selected and included in the GWAS Catalog. Finally, we used PLINK to extend the index SNPs on the 1000 Genomes Project phase 3 European data (window size = 1Mb, r^2^ > 0.8), and collectively defined the index SNPs and those in strong LD as the disease-related SNPs. A total of 159,358 neurological and 530,549 non-neurological disease-related SNPs were identified in different diseases (**Supplementary Table S10**).

### Variant functional annotation

All the 6,446,207 genotyped variants were functionally annotated using ANNOVAR software, a Perl command-line tool designed for efficiently and rapidly annotating the genomic variant data derived from high-throughput analysis (such as VCF file)^38^. The refGene database (hg38) was used as the annotation reference file consisting of functional information of variants in a genome-wide scale. It categorizes the variants as the following classes: ncRNA intronic, ncRNA exonic, ncRNA splicing, upstream, downstream, 5’-UTR, 3’-UTR, exonic, intronic, intergenic, and splicing site^61^. After removing 1,553 variants that are not annotated accurately (for example, annotated with multiple functional classes), we obtained a total of 6,444,654 variants annotated with unique functional classes. For the LD-pruned variants, the independent eSNPs and non-eSNPs were annotated into ten (lack of splicing variants) and 11 classes, respectively. Specifically, the numbers of the annotated independent eSNPs and non-eSNPs in the ten brain regions and whole brain were 646 and 592,299 (CRBL), 547 and 592,517 (FCTX), 470 and 595,412 (HIPP), 345 and 597,429 (MEDU), 460 and 593,566 (OCTX), 245 and 597,663 (PUTM), 167 and 596,922 (SNIG), 543 and 593,956 (TCTX), 294 and 596,187 (THAL), 549 and 590,478 (WHMT), and 2,628 and 1,927,132 (whole brain), respectively. The detailed statistics are described in Supplementary Table S16.

### Selection of TF binding sites

We downloaded the “brain-prefrontal-cortex 2022” (hg38) dataset was downloaded from a multiple-tissue TF ChIP-seq database ReMAP^41^. This dataset contains 133,084 peaks of TF SIN3A and OLIG2 binding sites in human FCTX, which is the only tissue that matched our study. Then, we selected the LD-pruned eSNPs and non-eSNPs of FCTX, and mapped them to the significant peak regions of the two TF binding sites by genomic coordinates, respectively.

### Correlation of lncRNA cis-eQTL effects among brain regions

Herein, we compared the similarity of genetic regulation on lncRNA expression between two of the ten brain regions by calculating the 𝑟_𝑏_ . It is an unbiased correlation estimate of eQTL effects, which accounts for both effect strength (beta values) and significance (errors or *P* values) in this process^43^. According to the previous study, we first selected the top-associated cis-eQTLs of the 425 elncRNAs (15 elncRNAs without eSNPs were removed) in each brain region, and extracted the eQTL beta coefficients, errors, and expression of the elncRNAs for subsequent analysis. Then, the Pearson correlation coefficient of each elncRNA expression level was calculated between two of the ten brain regions, and the correlation of errors in combination with the sample overlap was estimated based on the Bulik-Sullivan theory^62^. The variance of errors was approximated by the average of the square of the eQTL standard error. Using these statistics, we estimated the 𝑟_𝑏_ between the two brain regions by an extended Pearson correlation coefficient formula. Finally, the sampling variance of 𝑟_𝑏_ was calculated using the Jackknife approach, which excludes one elncRNA sequentially. See the original article for details on the algorithm^43^.

### Genetic prediction model training and TWAS

Firstly, we employed the PEER factors used for cis-eQTL analysis as covariates to conduct a multiple linear regression for each lncRNA and calculated the regression residuals to adjust the lncRNA expression matrices in each brain region. After integrating the adjusted lncRNA expression matrices, variant genotype data, and their corresponding annotation files (including IDs and locations of variants and lncRNAs), we trained the genetic prediction model in cis regions (within 1Mb of variant from lncRNA TSS) using the PredictDB pipeline that relies on an elastic net linear regression algorithm^63^. The trained model contains a database file and a SNP covariance matrices file. The database file undergoes further filtering based on the criterion of z-score P < 0.05 and average correlation coefficient (rho_avg) > 0.1. These two types of files were provided in lncBRAIN database. Finally, a large insomnia GWAS dataset comprising 1,331,010 individuals of European ancestry was utilized to perform TWAS analysis based on the trained model^64^. S-PrediXcan was employed to identify insomnia-related lncRNAs in each of the ten brain regions^65^.

### mRNA quantification and 2SLS estimation

Herein, we conducted a 2SLS estimation using eQTL data to avoid the impact of the potential unmeasured confounders while assessing the causal correlation between the exposure (lncRNA expression) and the outcome (mRNA expression). The LD-pruned variants significantly associated with lncRNA expression (eQTL nominal *P* < 10^-5^) are used as the instrumental variables. Firstly, the R package oligo with Affymetrix Exon 1.0 ST array annotation files was employed to quantify mRNAs from the same samples used for lncRNA quantification ^66^. Then, each instrumental variable was subjected to eQTL analysis with all the 17,008 quantified mRNAs, and the variants with smaller nominal *P* values in mRNAs than in lncRNAs were filtered out to satisfy the exclusion criteria. Subsequently, three distinct metrics were applied for quality control: (1) weak instrument test and Sargan test were used to examine whether the instrumental variables were sufficiently strong to ensure the validity of their estimates in the first stage (*P* < 0.05); (2) Wu-Hausman test was applied to verify the absence of potential endogeneity in the instrumental variable regression model, ensuring the robustness of the analysis (*P* > 0.05). Finally, the 2SLS estimation of each lncRNA with all mRNA individually was conducted using R package ivreg (https://CRAN.R-project.org/package=ivreg), and the *P* values were adjusted by the Benjamini-Hochberg method to identify the significant protein-coding genes whose expression is influenced by each lncRNA (with a threshold of FDR *q* < 0.05).

### Database construction for lncRNA expression regulation in human brain

The database lncBRAIN (http://lncbrain.org.cn/) visualizes the lncRNA expression, eQTL results, and the LD of the corresponding variants in the human brain. The results of lncRNA expression and eQTL analysis were visualized by box plots and distinguished according to gender, age, and brain region. The LD of variants in the cis-eQTL region was calculated and visualized using LDBlockShow software with its default parameters^67^. Users could easily query by lncRNA and SNP IDs, and download the results. The trained lncRNA prediction models and covariance matrices of the ten brain regions were also stored in lncBRAIN so users could to perform TWAS with their own data. We used HTML, CSS, and JavaScript to create interactive user interfaces, Python to handle back-end business logic and data flow, and MySQL to store the core data of the database.

## Acknowledgements

Authors would like to thank the United Kingdom Brain Expression Consortium (UKBEC) for sharing the dataset, and the Supercomputer Center of Chongqing Medical University for their computing power and technical support. This research is financially supported by the Science and Technology Research Program of National Natural Science Foundation of China (Grant No. 32200446), CQMU Program for Youth Innovation in Future Medicine, Chongqing Medical University (Grant No. W0147), and Chongqing Municipal Education Commission (Grant No. KJQN202100402).

## Data Availability

The datasets of lncRNA expression, cis and train-eQTL, trained lncRNA prediction models and covariance matrices for TWAS in the ten brain regions, and the LD results of variants in cis-eQTL region can be accessed and downloaded from the lncBRAIN database (http://lncbrain.org.cn/). Access to the other original data needs to be requested from the United Kingdom Brain Expression Consortium (UKBEC). The tools for quantifying lncRNA s by probe set re-annotation method are stored in the web server RElncANN (https://relncann.cqmu.edu.cn/appcel/). The codes used in this study are available at https://github.com/hyj260/LncBRAIN.

## Notes

### Competing Interest Statement

The authors have declared no competing interest.

