## Supplementary Figure S1 to S13 for "Genetic regulation of lncRNA expression in the whole human brain"

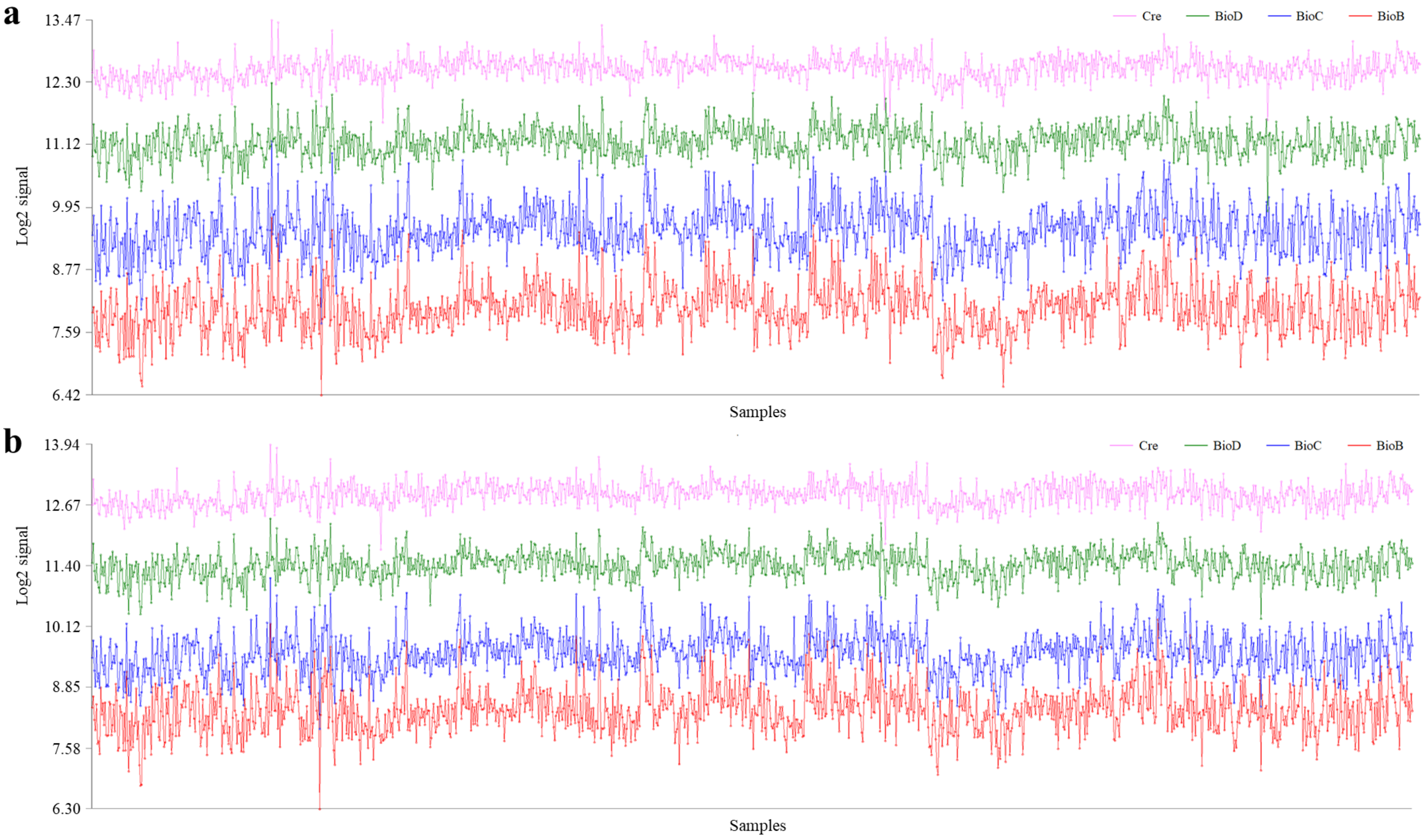


**Figure S1** Quality control of the Affymetrix Exon 1.0 ST arrays by the hybridization concentration assessment. The pre-labeled exogenous spike-in probes are BioB, BioC, BioD and Cre from 5' site (**a**) and 3' site (**b**), respectively.

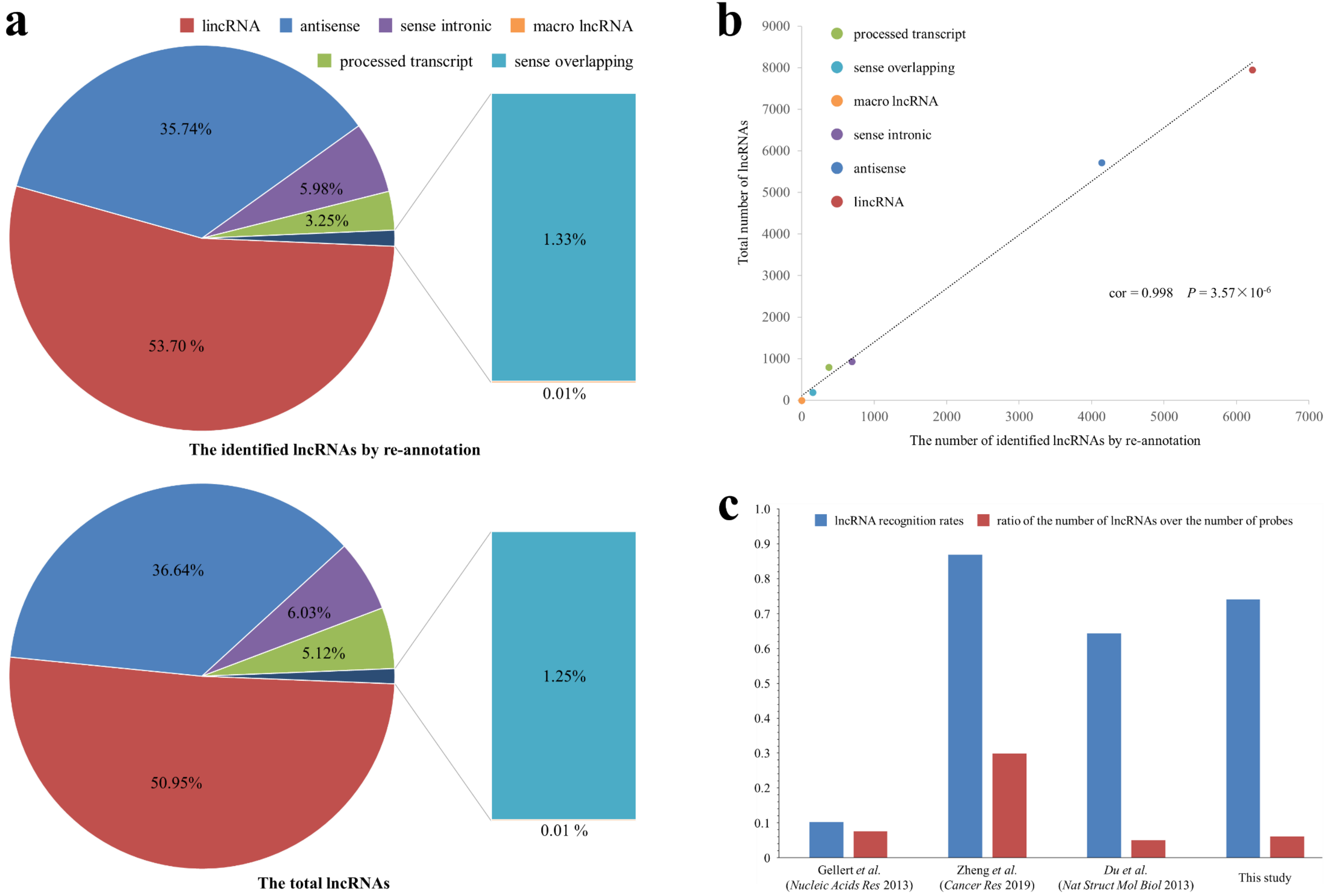
a
lincRNA
antisense
sense intronic
macro lncRNA
processed transcript
sense overlapping
35.74%
5.98%
3.25%
53.70 %
1.33%
0.01%
The identified lncRNAs by re-annotation
36.64%
6.03%
5.12%
50.95%
1.25%
0.01 %
The total lncRNAs
b
processed transcript
sense overlapping
macro lncRNA
sense intronic
antisense
lincRNA
Total number of lncRNAs
The number of identified lncRNAs by re-annotation
cor = 0.998 P = 3.57×10-6
0
1000
2000
3000
4000
5000
6000
7000
8000
9000
c
lncRNA recognition rates
ratio of the number of lncRNAs over the number of probes
1.0
0.9
0.8
0.7
0.6
0.5
0.4
0.3
0.2
0.1
Gellert et al. (Nucleic Acids Res 2013)
Zheng et al. (Cancer Res 2019)
Du et al. (Nat Struct Mol Biol 2013)
This study

**Figure S2** Affymetrix Exon 1.0 ST arrays probe set re-annotation for identifying lncRNAs. (**a**) Percentage of the six subtypes in the identified lncRNAs by re-annotation and the total lncRNAs, respectively. (**b**) Consistency of the proportions of the six subtypes in two groups. (**c**) Overall advantage of the improved probe set re-annotation approach used in this study compared with the previous methods.

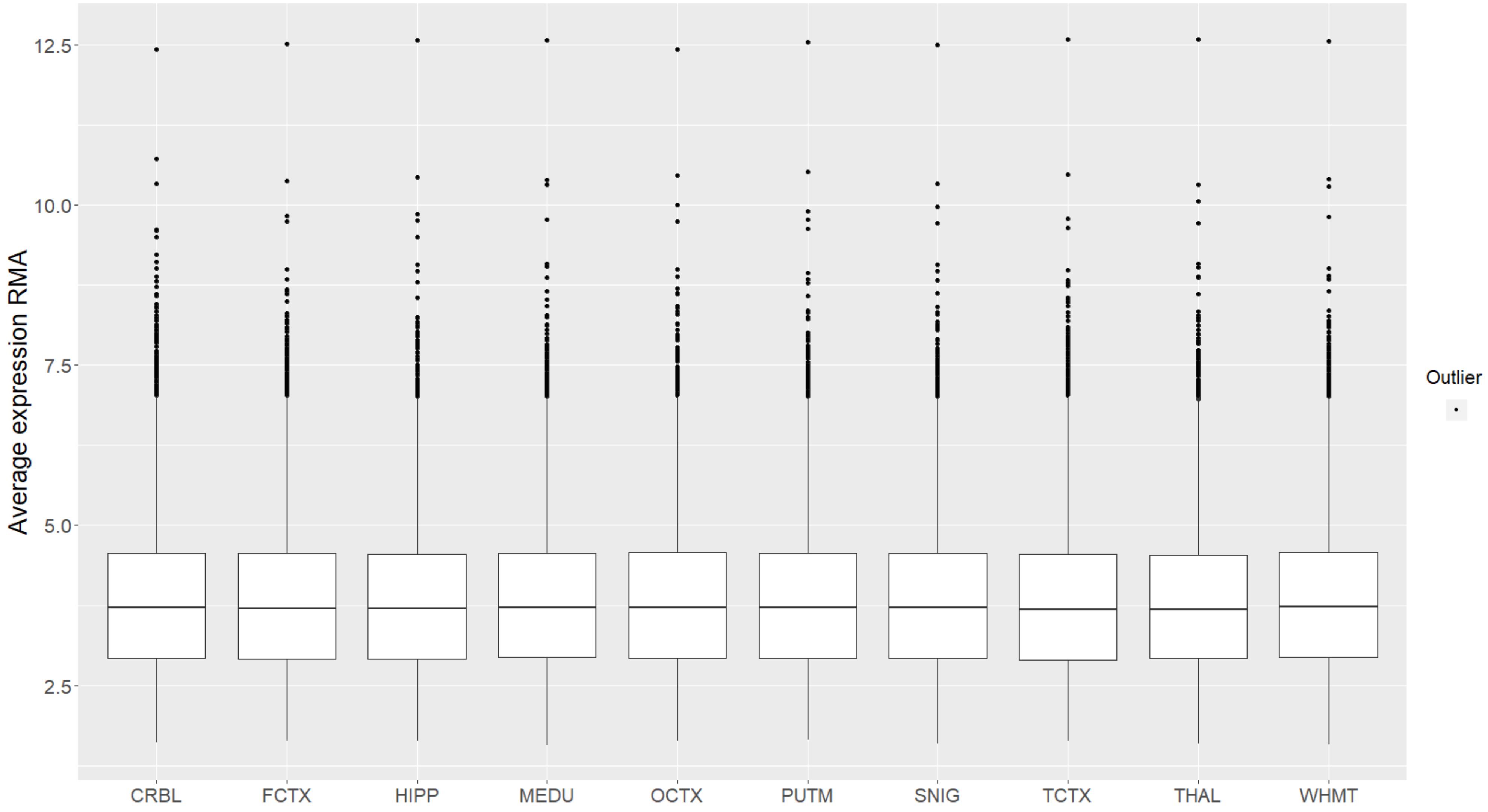


**Figure S3** The overall expression level of the 11,587 lncRNAs identified by exon arrays probe set re-annotation across the 10 brain regions. Each element represents the average expression of a lncRNA in a brain region.

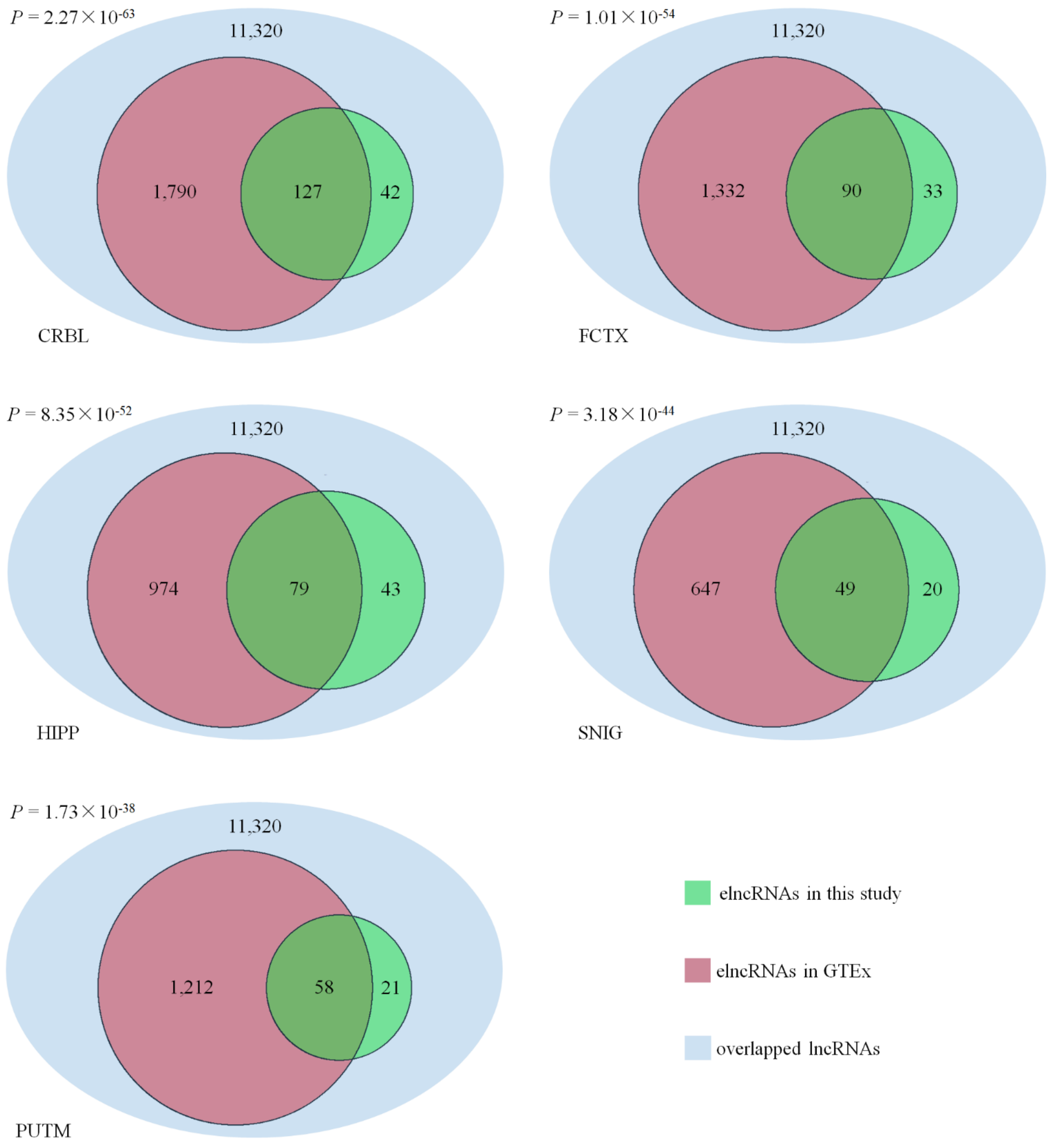


**Figure S4** The replication of our identified elncRNAs in brain region CRBL, FCTX, HIPP, SNIG, and PUTM in GTEx (v8) datasets. The statistical significance was determined by hypergeometric distribution test with the 11,320 overlapped lncRNAs as the background.

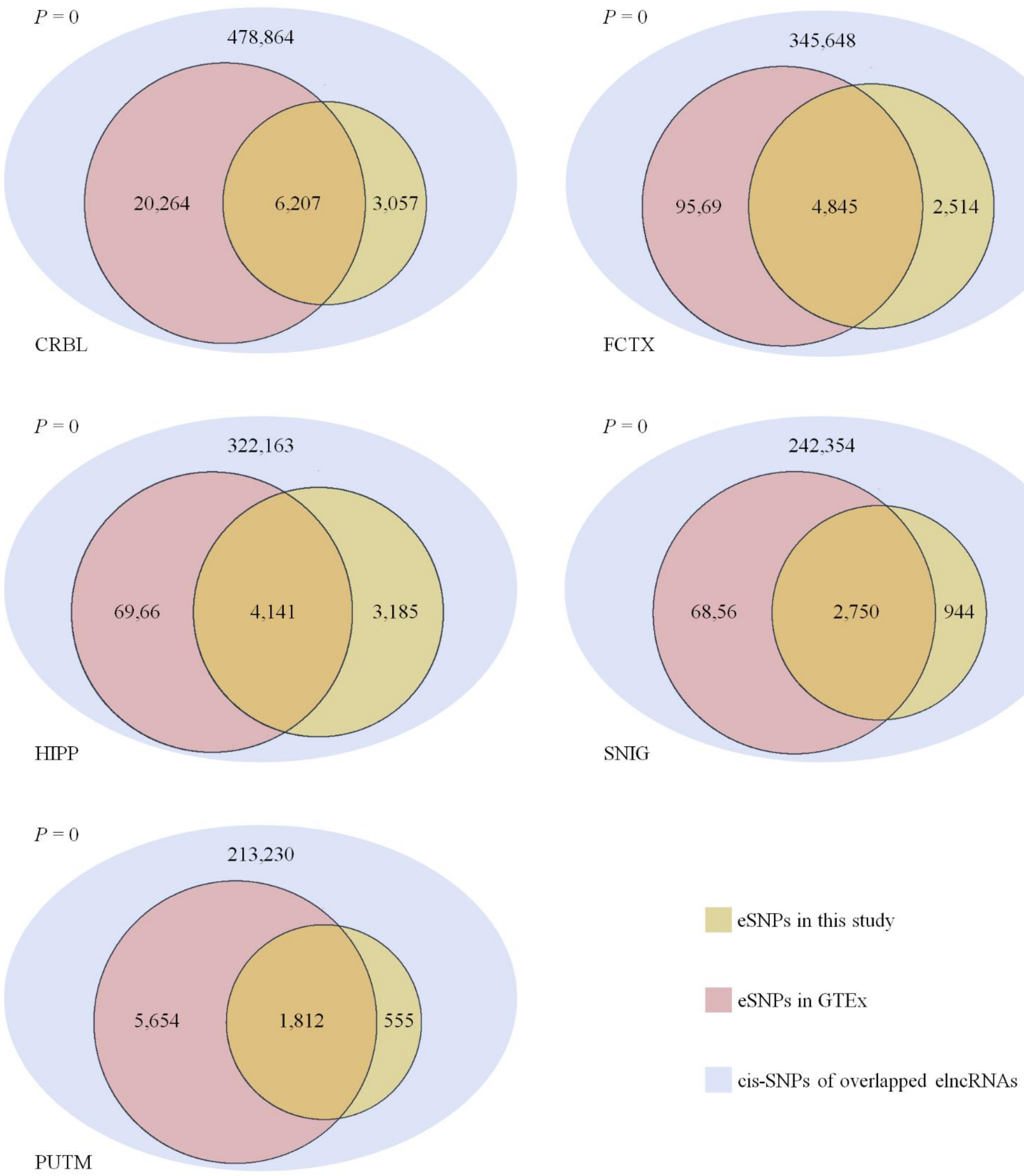


**Figure S5** The replication of our identified eSNPs in brain region CRBL, FCTX, HIPP, SNIG, and PUTM in GTEx (v8) datasets. The statistical significance was determined by hypergeometric distribution test with cis-SNPs of overlapped elncRNAs in each brain region as the background, respectively.

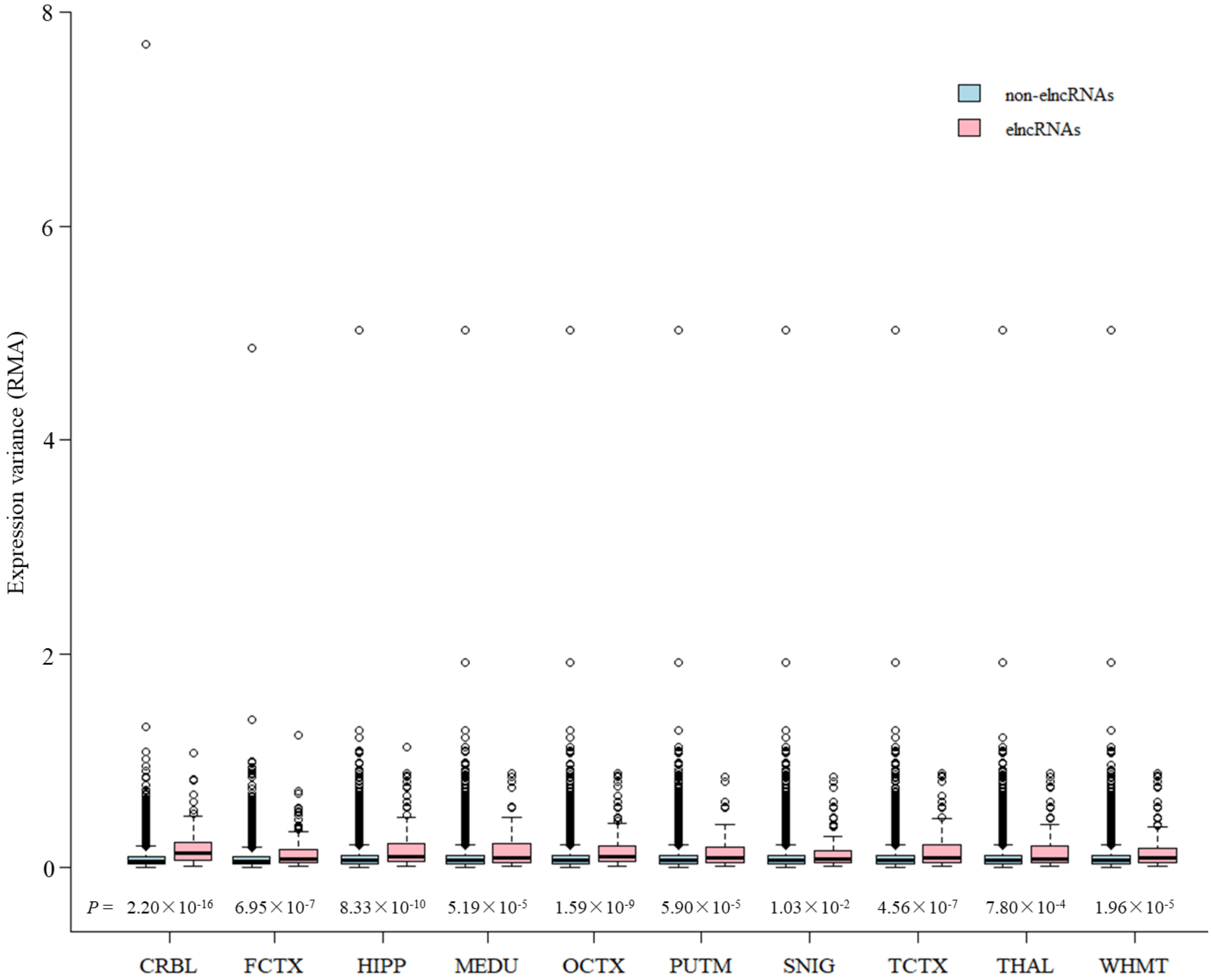


**Figure S6** The box plots show the expression variance of each elncRNA and non-elncRNA across individuals in the ten brain regions. The *P* values were calculated by two-tailed Wilcoxon test.

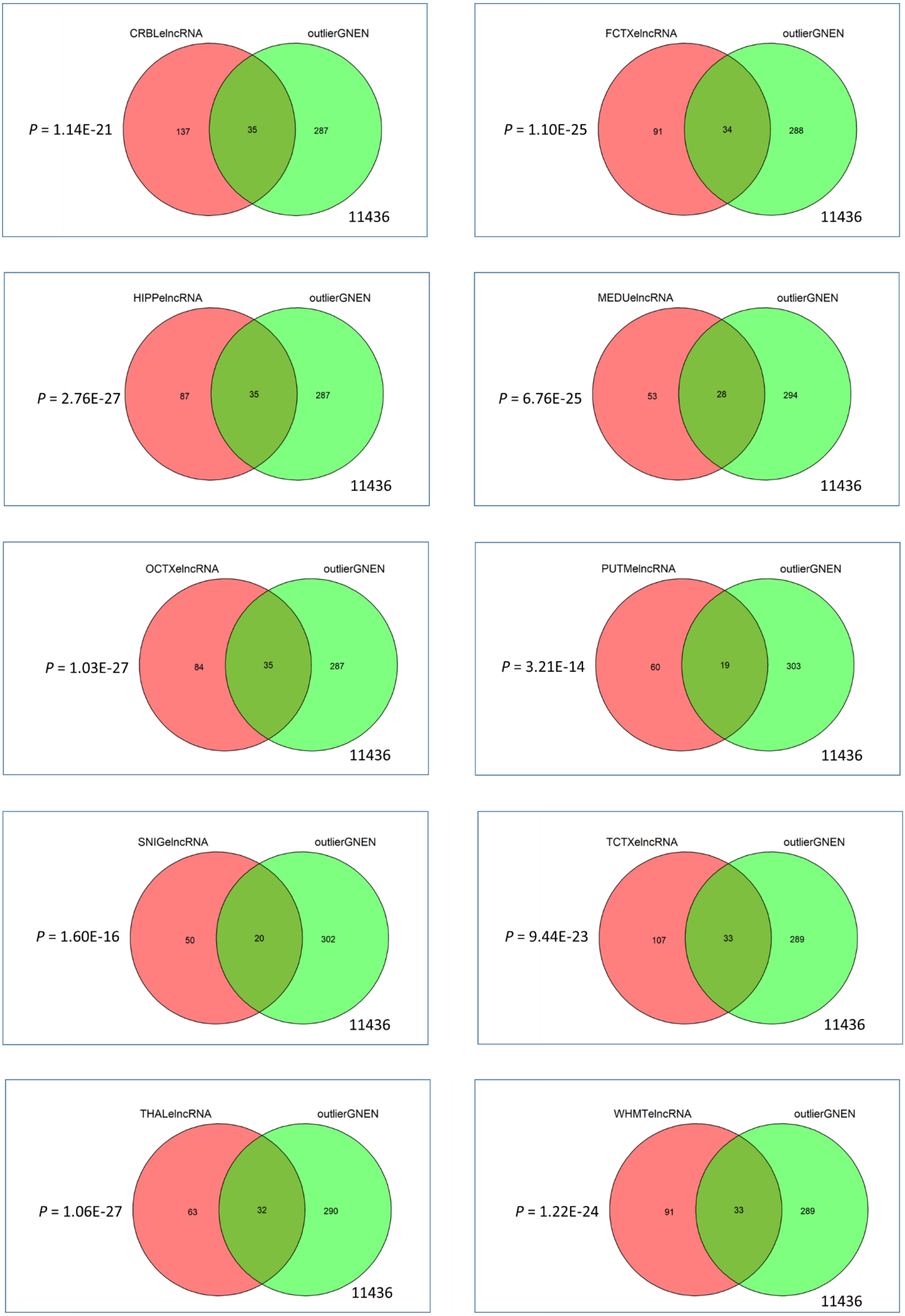
CRBLelncRNA
outlierGNEN
P = 1.14E-21
137
35
287
11436
FCTXelncRNA
outlierGNEN
P = 1.10E-25
91
34
288
11436
HIPPelncRNA
outlierGNEN
P = 2.76E-27
87
35
287
11436
MEDUelncRNA
outlierGNEN
P = 6.76E-25
53
28
294
11436
OCTXelncRNA
outlierGNEN
P = 1.03E-27
84
35
287
11436
PUTMelncRNA
outlierGNEN
P = 3.21E-14
60
19
303
11436
SNIGelncRNA
outlierGNEN
P = 1.60E-16
50
20
302
11436
TCTXelncRNA
outlierGNEN
P = 9.44E-23
107
33
289
11436
THALelncRNA
outlierGNEN
P = 1.06E-27
63
32
290
11436
WHMTelncRNA
outlierGNEN
P = 1.22E-24
91
33
289
11436

**Figure S7** The overlap between elncRNAs and outliers in each brain region. The statistical significance was determined by hypergeometric distribution test with the 11,436 lncRNAs used for eQTL analysis and outlier discovery as the background.

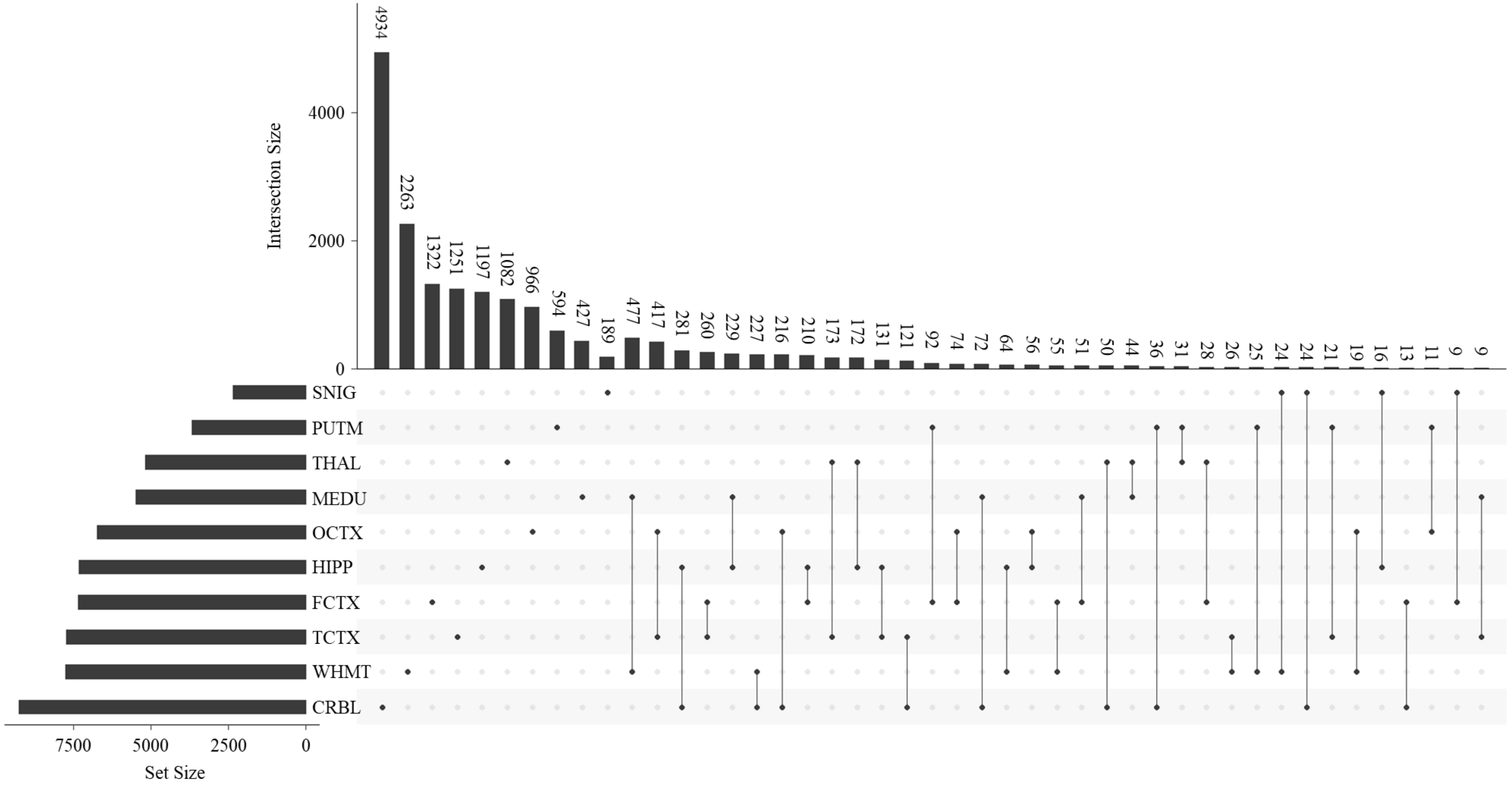


**Figure S8** The UpSet plot showing the number of eSNPs obtained from different brain regions and their tissue specificity and intersection.

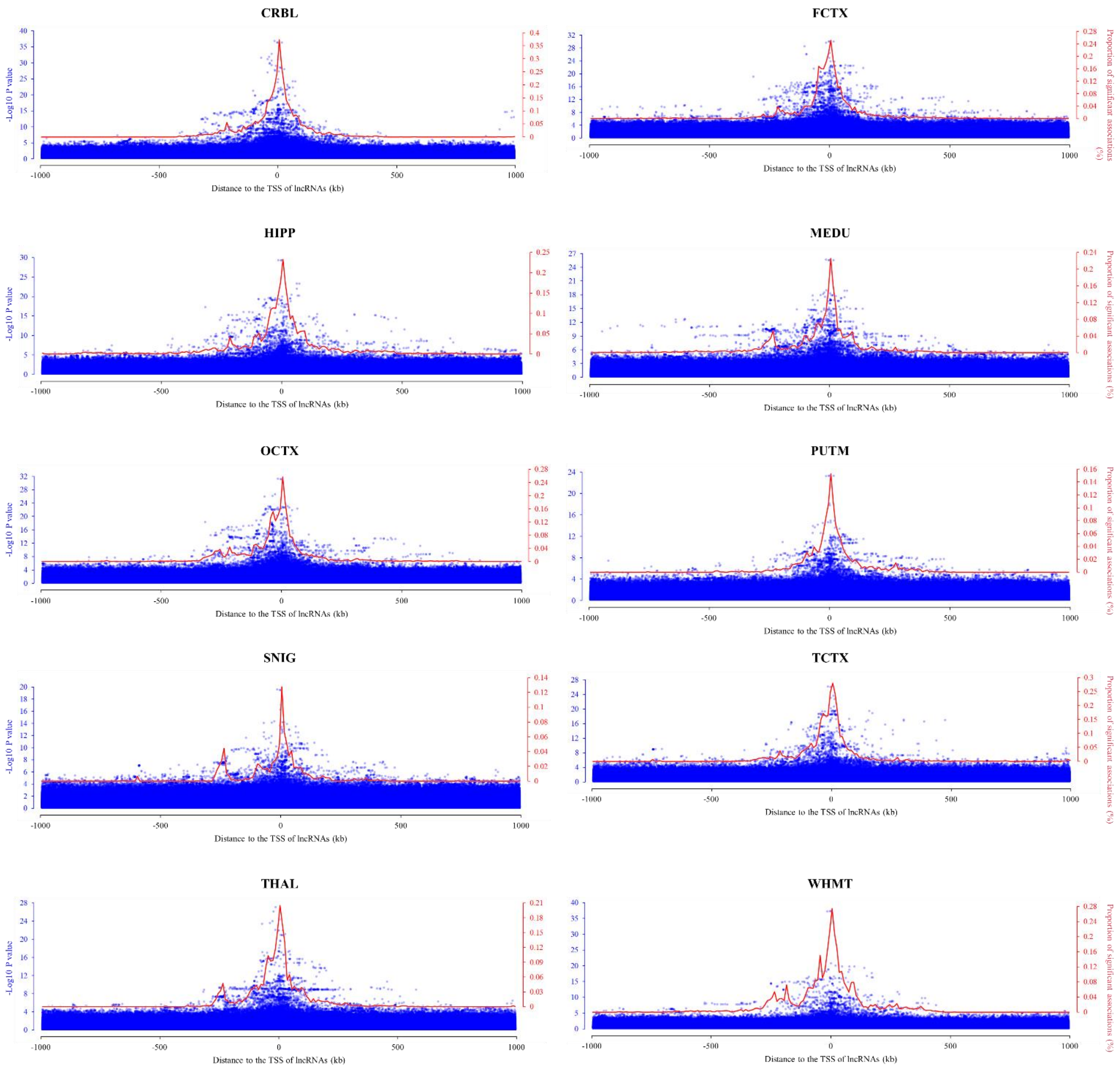


**Figure S9** Significance of eQTLs and percentage of eSNPs in relation to the distance from variants to TSS of corresponding lncRNAs in the ten brain regions. The blue dots represent the variants in cis-regions with the negative log-transformed eQTL P values (left y-axis). The red line represents the percentage of eSNPs per 10 kb window (right y-axis).

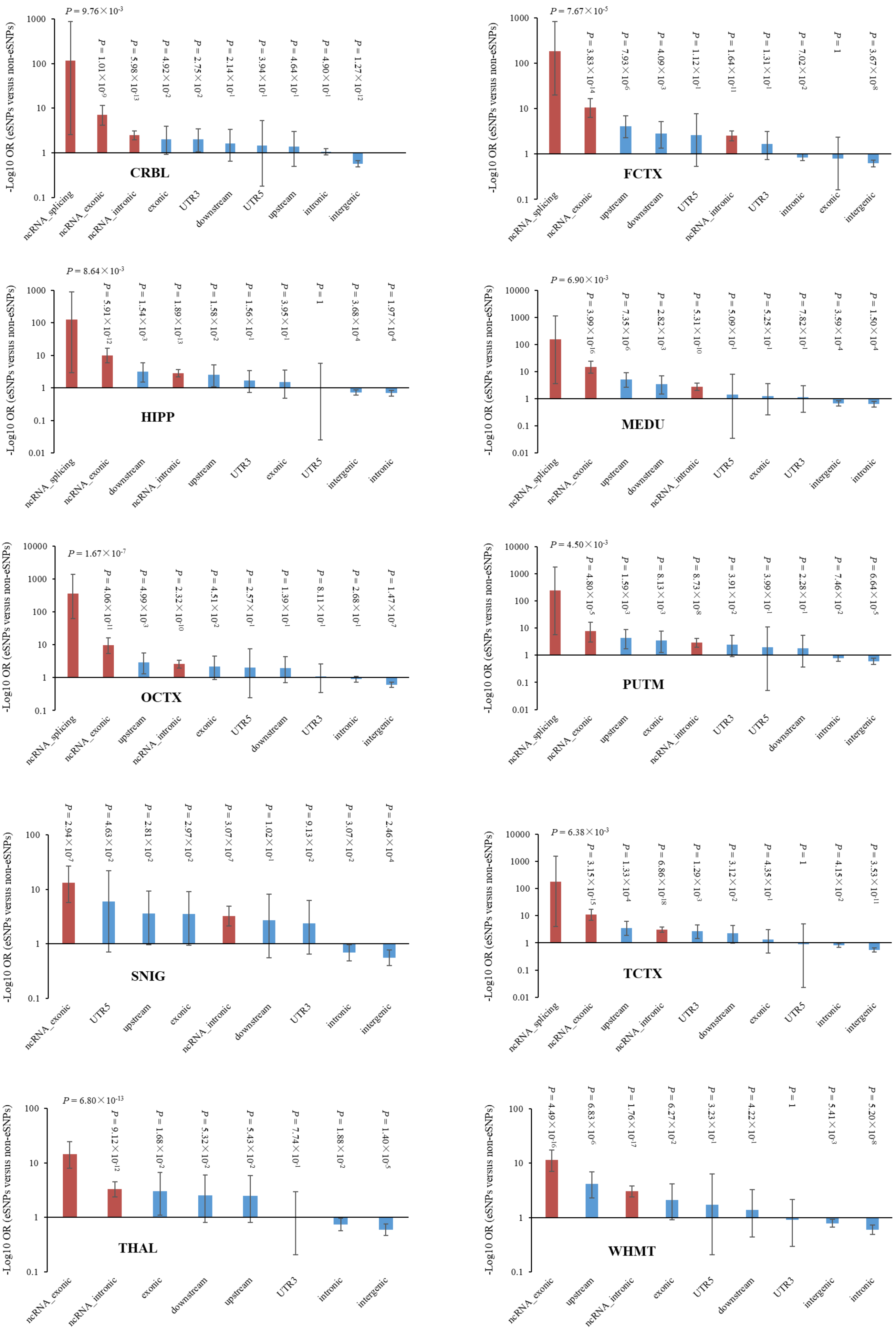

-Log10 OR (eSNPs versus non-eSNPs)
CRBL
$P = 9.76\times10^{-3}$
$P = 1.01\times10^{-9}$
$P = 5.98\times10^{-13}$
$P = 4.92\times10^{-2}$
$P = 2.75\times10^{-2}$
$P = 2.14\times10^{-1}$
$P = 3.94\times10^{-1}$
$P = 4.64\times10^{-1}$
$P = 4.90\times10^{-1}$
$P = 1.27\times10^{-12}$
ncRNA_splicing
ncRNA_exonic
ncRNA_intronic
exonic
UTR3
downstream
UTR5
upstream
intronic
intergenic
FCTX
$P = 7.67\times10^{-5}$
$P = 3.83\times10^{-14}$
$P = 7.93\times10^{-6}$
$P = 4.09\times10^{-3}$
$P = 1.12\times10^{-1}$
$P = 1.64\times10^{-11}$
$P = 1.31\times10^{-1}$
$P = 7.02\times10^{-2}$
$P = 1$
$P = 3.67\times10^{-8}$
ncRNA_splicing
ncRNA_exonic
upstream
downstream
UTR5
ncRNA_intronic
UTR3
intronic
exonic
intergenic
HIPP
$P = 8.64\times10^{-3}$
$P = 5.91\times10^{-12}$
$P = 1.54\times10^{-3}$
$P = 1.89\times10^{-13}$
$P = 1.58\times10^{-2}$
$P = 1.56\times10^{-1}$
$P = 3.95\times10^{-1}$
$P = 1$
$P = 3.68\times10^{-4}$
$P = 1.97\times10^{-4}$
ncRNA_splicing
ncRNA_exonic
downstream
ncRNA_intronic
upstream
UTR3
exonic
UTR5
intergenic
intronic
MEDU
$P = 6.90\times10^{-3}$
$P = 3.99\times10^{-16}$
$P = 7.35\times10^{-6}$
$P = 2.82\times10^{-3}$
$P = 5.31\times10^{-10}$
$P = 5.09\times10^{-1}$
$P = 5.25\times10^{-1}$
$P = 7.82\times10^{-1}$
$P = 3.59\times10^{-4}$
$P = 1.50\times10^{-4}$
ncRNA_splicing
ncRNA_exonic
upstream
downstream
ncRNA_intronic
UTR5
exonic
UTR3
intergenic
intronic
OCTX
$P = 1.67\times10^{-7}$
$P = 4.06\times10^{-11}$
$P = 4.99\times10^{-3}$
$P = 2.32\times10^{-10}$
$P = 4.51\times10^{-2}$
$P = 2.57\times10^{-1}$
$P = 1.39\times10^{-1}$
$P = 8.11\times10^{-1}$
$P = 2.68\times10^{-1}$
$P = 1.47\times10^{-7}$
ncRNA_splicing
ncRNA_exonic
upstream
ncRNA_intronic
exonic
UTR5
downstream
UTR3
intronic
intergenic
PUTM
$P = 4.50\times10^{-3}$
$P = 4.80\times10^{-5}$
$P = 1.59\times10^{-3}$
$P = 8.13\times10^{-3}$
$P = 8.73\times10^{-8}$
$P = 3.91\times10^{-2}$
$P = 3.99\times10^{-1}$
$P = 2.28\times10^{-1}$
$P = 7.46\times10^{-2}$
$P = 6.64\times10^{-5}$
ncRNA_splicing
ncRNA_exonic
upstream
exonic
ncRNA_intronic
UTR3
UTR5
downstream
intronic
intergenic
SNIG
$P = 2.94\times10^{-7}$
$P = 4.63\times10^{-2}$
$P = 2.81\times10^{-2}$
$P = 2.97\times10^{-2}$
$P = 3.07\times10^{-7}$
$P = 1.02\times10^{-1}$
$P = 9.13\times10^{-2}$
$P = 3.07\times10^{-2}$
$P = 2.46\times10^{-4}$
ncRNA_exonic
UTR5
upstream
exonic
ncRNA_intronic
downstream
UTR3
intronic
intergenic
TCTX
$P = 6.38\times10^{-3}$
$P = 3.15\times10^{-15}$
$P = 1.33\times10^{-4}$
$P = 6.86\times10^{-18}$
$P = 1.29\times10^{-3}$
$P = 3.12\times10^{-2}$
$P = 4.35\times10^{-1}$
$P = 1$
$P = 4.15\times10^{-2}$
$P = 3.53\times10^{-11}$
ncRNA_splicing
ncRNA_exonic
upstream
ncRNA_intronic
UTR3
downstream
exonic
UTR5
intronic
intergenic
THAL
$P = 6.80\times10^{-13}$
$P = 9.12\times10^{-12}$
$P = 1.68\times10^{-2}$
$P = 5.32\times10^{-2}$
$P = 5.43\times10^{-2}$
$P = 7.74\times10^{-1}$
$P = 1.88\times10^{-2}$
$P = 1.40\times10^{-5}$
ncRNA_exonic
ncRNA_intronic
exonic
downstream
upstream
UTR3
intronic
intergenic
WHMT
$P = 4.49\times10^{-16}$
$P = 6.83\times10^{-6}$
$P = 1.76\times10^{-17}$
$P = 6.27\times10^{-2}$
$P = 3.23\times10^{-1}$
$P = 4.22\times10^{-1}$
$P = 1$
$P = 5.41\times10^{-3}$
$P = 5.20\times10^{-8}$
ncRNA_exonic
upstream
ncRNA_intronic
exonic
UTR5
downstream
UTR3
intergenic
intronic

**Figure S10** Enrichment analysis of the eSNPs among different functional categories of variants in the ten brain regions measured by the two-tailed Fisher's exact test. The variants in non-coding genomic regions were marked red and others were marked blue. The black bars in histogram indicate 95% CI.

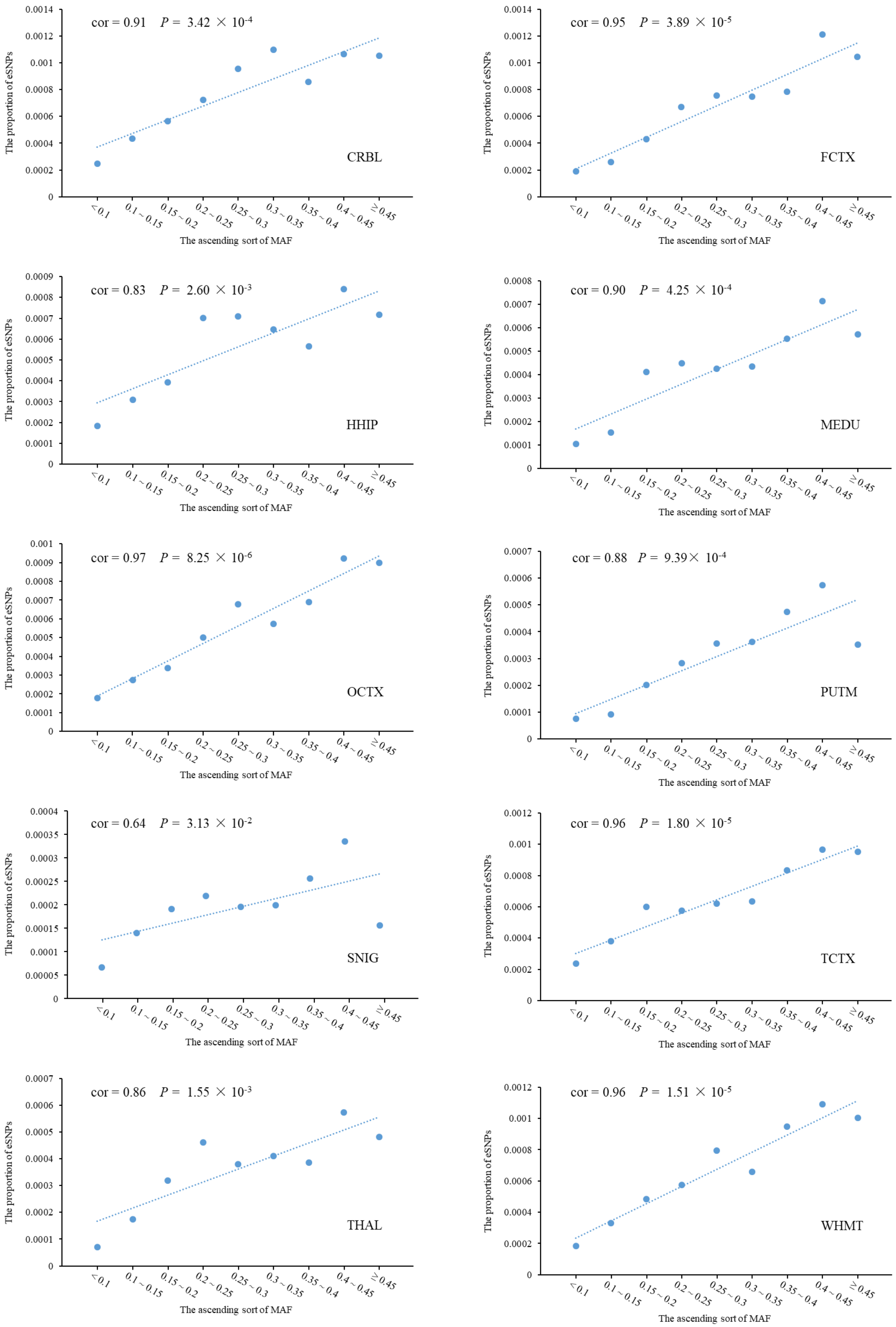
cor = 0.91 P = 3.42 × 10-4
CRBL
cor = 0.95 P = 3.89 × 10-5
FCTX
cor = 0.83 P = 2.60 × 10-3
HHIP
cor = 0.90 P = 4.25 × 10-4
MEDU
cor = 0.97 P = 8.25 × 10-6
OCTX
cor = 0.88 P = 9.39× 10-4
PUTM
cor = 0.64 P = 3.13 × 10-2
SNIG
cor = 0.96 P = 1.80 × 10-5
TCTX
cor = 0.86 P = 1.55 × 10-3
THAL
cor = 0.96 P = 1.51 × 10-5
WHMT
The proportion of eSNPs
The ascending sort of MAF
<0.1
0.1～0.15
0.15～0.2
0.2～0.25
0.25～0.3
0.3～0.35
0.35～0.4
0.4～0.45
≥0.45

**Figure S11** The change in proportion of eSNPs among cis-SNPs with increasing MAF in the ten brain regions. The blue points represent the proportions in corresponding MAF bin, and the dotted lines are their linear regression results.

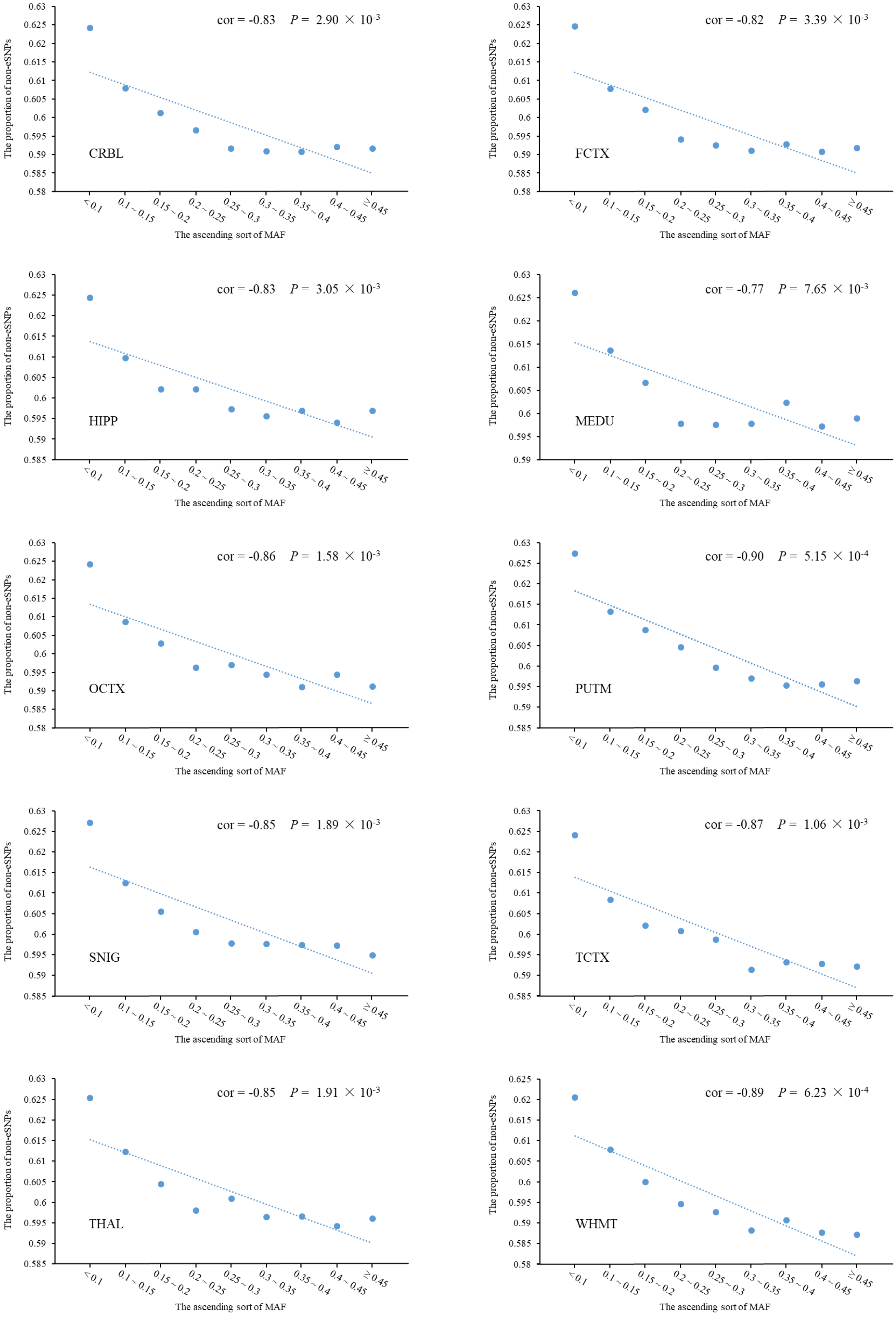

cor = -0.83 P = 2.90 × 10-3
CRBL
cor = -0.82 P = 3.39 × 10-3
FCTX
cor = -0.83 P = 3.05 × 10-3
HIPP
cor = -0.77 P = 7.65 × 10-3
MEDU
cor = -0.86 P = 1.58 × 10-3
OCTX
cor = -0.90 P = 5.15 × 10-4
PUTM
cor = -0.85 P = 1.89 × 10-3
SNIG
cor = -0.87 P = 1.06 × 10-3
TCTX
cor = -0.85 P = 1.91 × 10-3
THAL
cor = -0.89 P = 6.23 × 10-4
WHMT
The proportion of non-eSNPs
The ascending sort of MAF
<0.1
0.1~0.15
0.15~0.2
0.2~0.25
0.25~0.3
0.3~0.35
0.35~0.4
0.4~0.45
≥0.45

**Figure S12** The change in proportion of non-eSNPs among cis-SNPs with increasing MAF in the ten brain regions. The blue points represent the proportions in corresponding MAF bin, and the dotted lines are their linear regression results.

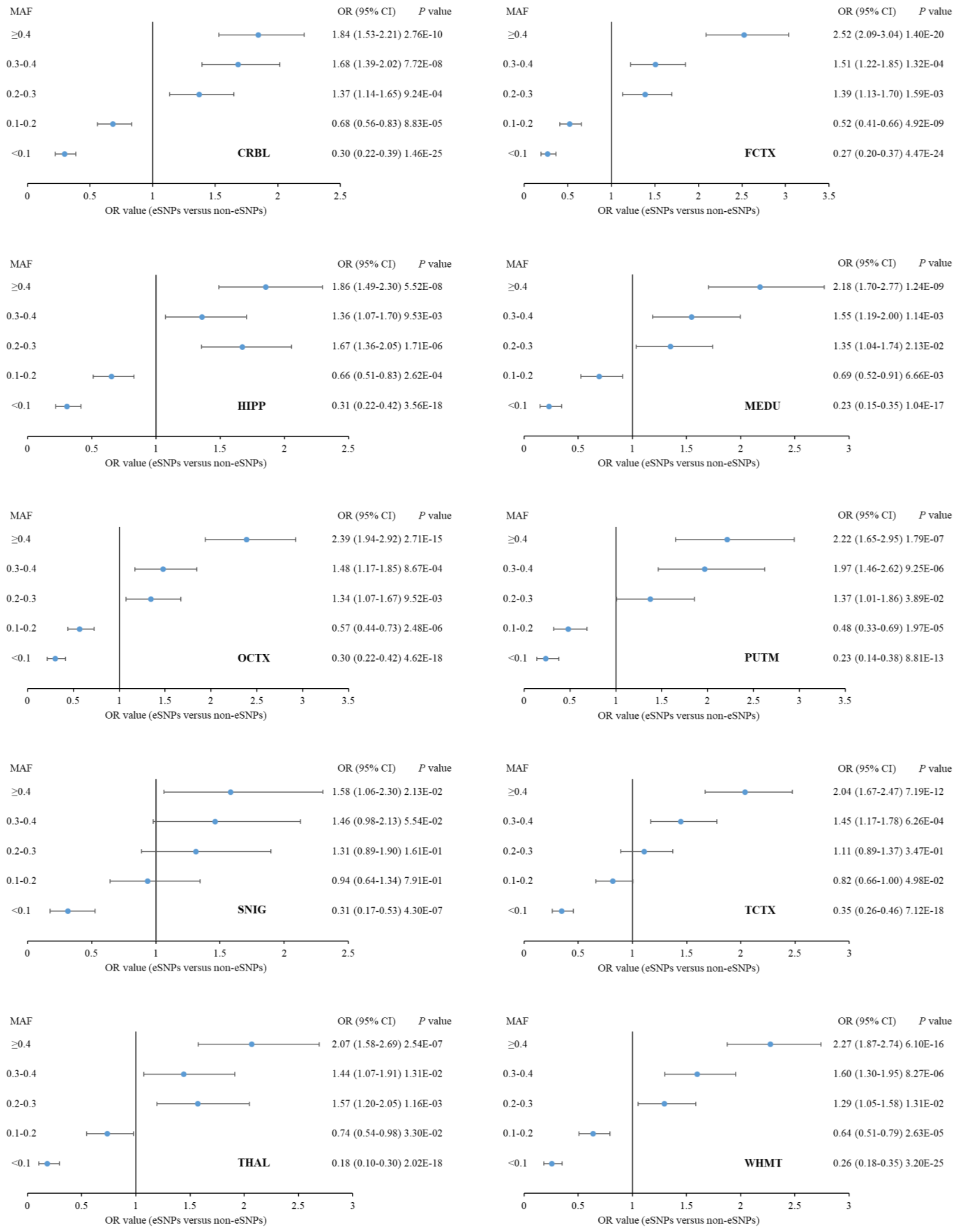


**Figure S13** The forest plot showing the enrichment results of eSNPs when compared with the non-eSNPs in each MAF bin of each brain region. The OR and 95% CI were calculated by two-tailed Fisher's exact test.
